# PAMP-Induced secreted Peptide-Like 6 (PIPL6) functions as an amplifier of plant immune response through RLK7 and WRKY33 module

**DOI:** 10.1101/2022.11.30.518506

**Authors:** Javad Najafi, Ragnhild Sødal Gjennestad, Ralph Kissen, Tore Brembu, Zdenka Bartosova, Per Winge, Atle M Bones

## Abstract

Plant peptide hormones are engaged in the regulation of plant developmental programs and immunity. PAMP-Induced Peptide (PIP) hormones are new class of signaling peptide with diverse functional roles in the regulation of plant development and stress responses. In this study, we have investigated the function of PAMP-Induced secreted Peptide-Like 6 (PIPL6) as an amplifier of plant immunity against necrotrophic fungal pathogens in *Arabidopsis thaliana*. We have applied an integrated omics approach to unveil the function and downstream signaling pathways initiated by PIPL6. *PIPL6* is highly and transiently induced by treatment with different elicitors. Exogenous application of synthetic peptide designed from the C-terminal conserved domain of PIPL6 resulted in strong transcriptional induction of many genes involved in the regulation of plant immunity. Further gene expression analysis revealed that induction of marker genes by PIPL6 peptide requires the receptor-like kinase 7 (RLK7). Immunoblotting and gene expression analysis demonstrated that exogenous applications of PIPL6 peptide activates MAPK6, MAPK3, and WRKY33 module in an RLK7-dependent manner. The levels of salicylic acid, jasmonic acid, camalexin, and glucosinolates were differentially regulated in *PIPL6* knock-down and overexpression lines challenged by necrotrophic pathogen *Botrytis cinerea*. Bioassays using the necrotrophic fungal pathogens *Botrytis cinerea* and *Alternaria brassicae* showed that *pipl6* knock-down lines were more susceptible to these pathogens while *PIPL6* overexpression lines exhibited enhanced resistance. Altogether, these results indicate that the PIPL6 peptide functions as a new damage-associated molecular pattern (DAMP) and acts as an amplifier of Arabidopsis immunity.

## Introduction

As sessile organisms, plants are highly vulnerable to be attacked by a diverse range of microbial pathogens, viruses, insects and other herbivores. In response, plants have evolved to mount sophisticated defense mechanisms to defend themselves against attackers. The first line of defense against the invading pathogens depends on receptor proteins localized in the cell membrane termed as pattern recognition receptors (PRRs) that recognize pathogen-associated molecular patterns (PAMPs) or microbe-associated molecular patterns (MAMPs) (Jones and Dangl, 2006). PAMPs/MAMPs are conserved, microbial-specific molecules from pathogens, such as flagellin, lipopolysaccharides, and chitin. Pattern recognition receptor (PRR) proteins have a conserved structure consisting of an extracellular PAMP interacting domain and in many cases harboring an intracellular protein kinase domain. Several plant PRRs have been characterized, including the receptors for bacterial flagellin (FLS2) (Gómez-Gómez and Boller, 2000) and EF-Tu (EFR) (Zipfel et al., 2006), and fungal chitin (CERK1) (Miya et al., 2007). Perception of DAMPs/MAMPs by their corresponding receptors is followed by a burst of Ca^2+^ (Hu et al., 2004) and reactive oxygen species (ROS) (Miller et al., 2009), activation of mitogen-activated protein (MAP) kinase signaling cascades (Ligterink et al., 1997), transcriptional reprogramming (Maleck et al., 2000), deposition of callose at the site of infection (Luna et al., 2011), and production of phytoalexins (Ahuja et al., 2012) and other secondary metabolites involved in plant defense responses (Bednarek, 2012).

Plant immune responses can also be induced by endogenous molecules produced upon pathogen attack such as cell-wall fragments and peptides. Small signalling peptides have attracted attention for their role in the regulation of plant immunity (Boller and Flury, 2012). AtPEP1 was the first peptide with damage-associated molecular pattern (DAMP) activity purified from Arabidopsis (Huffaker et al., 2006) that is perceived by two receptor-like kinases, PEP RECEPTOR 1 (PEPR1) (Krol et al., 2010) and PEPR2 (Yamaguchi et al., 2010). Perception of PEP1 by PEPR1 and PEPR2 receptors enhances host plant resistance against the biotrophic pathogens *Pseudomonas syringae* and *Pythium irregulare* and the necrotrophic pathogen *Botrytis cinerea* through activation of BOTRYTIS-INDUCED KINASE 1 (BIK1) (Liu et al., 2013). From a structural point of view, the PEP1 precursor lacks an N-terminal signal peptide that would direct it to the apoplast. Hence, it is suggested that physical damage caused by pathogen attack or wounding facilitates PEP1 translocation to the apoplast where it interacts with its receptors, initiates danger/alarm signaling pathways and couples local and systemic pathways to amplify immune responses (Ross et al., 2014).

Both local and systemic responses are regulated by phytohormone initiated pathways and their cross-talk depending on the nature of the invaders and their lifestyle (Spoel and Dong, 2008). Among phytohormones salicylic acid (SA), jasmonic acid (JA), ethylene (ET), and abscisic acid (ABA) are studied more comprehensively for their functions in the regulation of plant immune responses. SA mediates plant response against biotrophs, whereas JA and ET cooperatively regulate plant immunity to mount proper responses against necrotrophic pathogens and herbivorous insects (Glazebrook, 2005). Triggering of immune related phytohormones biosynthesis requires changes in gene expression including those encoding transcription factors (TFs). The WRKY family of TFs have been comprehensively investigated for their roles in the regulation of plant defence responses. It has been shown that WRKY33 plays a vital role in the biosynthesis of salicylic acid (SA), jasmonic acid (JA), and camalexin when Arabidopsis is infected with *B. cinerea* (Birkenbihl et al., 2012).

In addition to the PEPs, members of another family of plant endogenous signalling peptides termed PAMP-induced secreted peptides (PIPs) with DAMP activity have been characterized in Arabidopsis. It has been shown that PIP1 and PIP2 amplify immunity through RECEPTOR-LIKE KINASE 7 (RLK7) against *Pseudomonas syringae* and *Fusarium oxysporum* (Hou et al., 2014). An opposite role was reported for PIP3 and detailed studies showed that plants overexpressing *prePIP3* are susceptible against both biotrophic and necrotrophic pathogens (Najafi et al., 2020).

Here, we investigated the function of PAMP-induced secreted peptide like 6 (PIPL6) in the regulation of plant immunity against necrotrophic fungal pathogens in *Arabidopsis thaliana*. Treatment of wildtype (Wt) plants with flagellin22 (flg22) and PEP1 peptides, and chitin hexamers revealed that *PIPL6* expression is highly and transiently induced after applied elicitors. Ectopic application of synthetic PIPL6 peptide followed by total transcriptomic analysis using RNA-seq showed that expression of many immune and detoxification related genes are highly induced. Immunoblotting assays showed activation of the MAP kinase (MPK3/MPK6) and WRKY33 module upon PIPL6 treatment. Both transcriptome reprogramming, MPK3/6 phosphorylation, and WRKY33 induction upon PIPL6 application require RLK7 as a receptor. Analysis of PIPL6 transgenic knock-down and overexpression lines revealed altered phenotype and profiles of phytohormones, camalexin, and glucosinolates after necrotrophic fungal pathogen *Botrytis cinerea* treatment. These findings suggest that PIPL6 functions as a DAMP peptide and amplifies plant immune responses.

## Materials and Methods

### Plant materials and growth conditions

*Arabidopsis thaliana* ecotype Columbia-0 (Col-0) was used as the wild-type (Wt.) control. All mutants and transgenic plants used in this study are in the Col-0 background. Seeds were surface sterilized and grown under long days condition (16 hours light (150 *μ*mo1 m^-2^s^-1^), 8 hours dark at 22°C 20-25% relative humidity. Two mutant lines for *PIPL6* were screened, *pipl6-1* (SALK_106769) that has a T-DNA inserted in the promotor region, and *pipl6-2* (Wisc_DsLoxHs144_04E.1) with a transposon inserted immediately after the stop codon of the *PIPL6* coding sequence. Homozygous plants were screened based on growth on selection medium and PCR using a combination of gene-specific and transposon or T-DNA-specific primers (Supplementary Table 1). Other mutant lines used in this study including *rlk7, wrky18, wrky60, wrky40* and double mutants *wrky18/40, wrky40/60, wrky18/60* were described previously by Najafi *et al*. (Najafi et al., 2020). *Wrky33* and WRKY:HA line expressing a *proWRKY33*:WRKY33:HA construct in this *wrky33* background were described by Birkenbihl *et al*. (Birkenbihl et al., 2012)

Overexpression lines of *PIPL6* (At1g47178) were generated by PCR amplification of the coding sequence from Col-0 plants and subsequent cloning into the destination vector pEG100 (Earley et al., 2006) under control of the 35S promoter using Gateway^®^ technology. The construct was introduced to *Agrobacterium tumefaciens* strain C58C1 pGV2260 and transformed into Col-0 Wt plants using the floral dip method (Clough and Bent, 1998). Two Independent transgenic T3 lines with a single copy of T-DNA and constitutive expression of *PIPL6* were selected for further analysis.

### Elicitor and peptide treatments

flg22 (QRLSTGSRINSAKDDAAGLQIA), PEP1 (ATKVKAKQRGKEKVSSGRPGQHN) and chitin hexamers were used as elicitors to treat wild-type plants followed by *PIPL6* gene expression analysis. Wild-type seeds were grown at the density of 10-12 seeds per well in 6-well culture plates containing 1,5 mL of 1/2MS liquid medium supplemented by 1% sucrose per well in three biological replicates (3 wells) for each timepoint. The seedlings were grown for 7 days under long day conditions (16h/8h). At day eight, treatments were performed by replacing the growth medium with fresh medium containing flg22 and PEP1 at the final concentrations of (100 nM) and chitin hexamer (500 nM). Synthetic PIPL6 was designed from the C-terminus of the prePIPL6 peptide containing the conserved SGPS motif with the following sequence: (AFRLA**SGPS**RKGRGH). The effects of synthetic PIPL6 peptide on the expression of immune marker genes and total transcriptome profiling were examined in both wild-type and *rlk7*. Peptides were synthesized with a purity of >95% by Biomatik (Cambridge, Ontario, Canada). PIPL6 peptide was sprayed on two-week-old seedlings grown on solid 1/2MS medium supplemented by 1% sucrose at the final concentration of 1 μM as described by previously (Najafi et al., 2020). Three hours after treatment, rosette leaves were harvested in four biological replicates. Each replicate consisted of material pooled from four petri dishes.

### Gene expression analysis (qRT-PCR and RNA-seq)

Harvested tissues from different treatments were subjected for RNA isolation, cDNA synthesis, and qRT-PCR as described by Vie et al.(Vie et al., 2015). For RNA-seq analysis, three replicates for control and PIPL6 peptide treatment of wild-type seedlings were sampled three hours after treatment. RNA integrity was examined on Aligent 2100 Bioanalyzer using the Aligent RNA 6000 Nano kit (Aligient Technologies). RNA sequencing was performed by GENEWIZ (Leipzig, Germany) on an Illumina sequencing platform. The raw sequence reads in FASTq format were mapped against Arabidopsis TAIR10 gene models with the Bowtie2 sequence mapper in a very-sensitive-local model (Langmead and Salzberg, 2012). The sequence Alignment/Map file generated by the Bowtie2 was used to produce a “count-table” were reads mapping specific Arabidopsis genes were registered for each of the individual biological replicates. The edgeR software package was used for statistical analysis of the mapped sequence reads in the count table (Robinson et al., 2010). Genes with low expression were identified and filtered out, according to the following criteria: a gene cut-off at 1 hit per 1 million reads, keeping only genes that were above the cut-off for all biological replicates. The data was analysed using statistical methods based on generalized linear models, and a likelihood ratio test was used to identify differentially expressed genes. Genes with a false discovery rate below 0.01 and log2 value > ± 1 were defined as significantly differentially expressed genes. Gene ontology and data enrichment analysis was performed in Cytoscape environment and ClueGo plug-in (Bindea et al., 2009).

### Growth inhibitory and ROS assays

Growth inhibition in response to the elicitor flg22 was performed for Wt and *PIPL6* knock-down and overexpression lines. Seeds were grown on 1/2MS agar plates supplemented by 1% sucrose under long day conditions for five days. At day five, seedlings were transferred to a 48-well-plate with 1/2MS liquid medium supplemented by 1% sucrose (one seedling per well with 600μL medium) containing flg22 (100nM) or only liquid medium as a control. For each genotype 24 seedlings for control treatment and 48 seedlings for flg22 treatment were used. The plates were incubated under long day conditions for the next 10 days and fresh biomass was measured. The flagellin insensitive *fls2* line was used as a control in the assay.

For the ROS assay, plants were grown under short day conditions (10/14 light-dark photoperiod) for 4-5 weeks prior to experiments. Luminol-based ROS generation detection was performed as described by Bisceglia *et al* (Bisceglia et al., 2015).

### Immunoblotting

For the MAP kinase activation assay, Wt and *rlk7* seeds were grown in 1ml of 1/2MS liquid medium supplemented by 0.25% sucrose in a 12-well culture plate for ten days at a density of 10 seeds per well. One day before treatments, the culture was replaced with fresh medium. Seedlings were treated either with 1 μM of flg22 or PIPL6 synthetic peptides with water as control. Samples were harvested at defined time points, flash frozen in liquid N2 and stored at −80°C for protein extraction with the buffer (25 mM Tris-HCl pH7.8, 75 mM NaCl, 15 mM EGTA, 10 mM MgCl2, 15 mM B-Glycerophosphate, 15 mM 4-Nitrophenylphosphate bis, 1 mM DTT, 1mM NaF, 0.5 mM Na3VO4, 0.5 mM PMSF, 1% Protease inhibitor cocktail (Sigma, P9599) and 0.1% Tween20). 40 μg of total proteins were separated in 10% SDS-PAGE, transferred to nitrocellulose membrane, probed overnight using Phospho-p44/p42 MAPK (Erk ½)(Thr202/Tyr204) antibody (4370; Cell Signaling Technology), and incubated with goat antirabbit horseradish peroxidase (HRP)-conjugated secondary antibody (A0545, Sigma-Aldrich). HRP activity was developed with a ECL chemiluminescent reagent (Bio-Rad) and detected using a G:Box XRQ imaging system (Syngene). For WRKY33 activation, transgenic seeds expressing a *proWRKY33*:WRKY33:HA construct in the *wrky33* background were grown and treated under the same conditions as explained above. Proteins were isolated from samples harvested at different time points and 5μg of total proteins were separated in 10% SDS-PAGE, transferred to nitrocellulose membrane. Blots were incubated over-night in Anti-HA antibody (SantaCruz, SC-805), followed by goat anti-rabbit horseradish peroxidase (HRP)-conjugated secondary antibody (A0545, Sigma-Aldrich), an ECL chemiluminescent reagent (Bio-Rad) and detected using a G:Box XRQ imaging system (Syngene) to probe for WRKY33 induction and abundance. Experiments were repeated two times with similar responses.

### Pathogen assay and metabolite analysis

Wild-type, *wrky33*, PIPL6 knock-down and overexpression lines were grown for five weeks under short days (10/14 hours photoperiod) condition. Plants were challenged either by *Botrytis cinerea* isolate 2100 (CECT2100; Spanish type) spores or Vogel buffer as control as described by Birkenbihl et al.(Birkenbihl et al., 2012). Phenotype of plants were assessed five days post infection (dpi). For metabolite analyses, plants were sprayed using fungal spores to a density of 2.5×10^5^ spores ml^-1^. Rosette leaves collected 48 hours post infection (hpi) in five biological replicates and flash frozen in liquid N2 and stored at −80°C for further applications. Each replicate was defined as a pot containing three plants. Tissues from individual replicates were divided into three parts. One part was used for RNA extraction and gene expression analysis by qRT-PCR as previously described. The second part was used for phytohormone and camalexin extraction. The last part was used for glucosinolate extraction. Phytohormones and camalexin were extracted following the method described by Salme et al. (Salem et al., 2020) Briefly, 100 mg of frozen tissue was subjected for phytohormone extraction using 1 ml of pre-cooled (−20°C) extraction solvent tert-Butyl methyl ether (MTBE):methanol (3:1 V/V) spiked by deuterated JA+SA+ABA at final concentrations of 5+5+1 ng/ml as internal standards. For camalexin, pure camalexin (SML1016, Sigma-Aldrich) dissolved in ethanol was used as external standard in five replicates. Samples were vortexed and kept on an orbital shaker for 30 min at 4°C followed by a 15min sonication step in an ice-cooled sonicator bath. A volume of 0.5 ml of acidified water (0.1% HCl) was added and the samples were vortexed for 1 min. Samples were kept on an orbital shaker for an additional 30 min at 4°C and centrifuged at 10,000 *g* for 10 min at 4°C. A fixed volume (0.7 ml) of the upper supernatant (MTBE phase) was transferred to a fresh 1.5 ml microcentrifuge tube and dried using a Speed-Vac concentrator at room temperature. The dried pellets were resuspended in 100 μl of water:methanol (1:1 V/V) solvent and the resuspended samples were immediately subjected to UPLC-ESI-MS/MS analysis.

Glucosinolates were analysed as described by Kissen et al.(Kissen et al., 2016). Briefly, 100 mg of plant tissue was freeze-dried and glucosinolates were extracted in boiling methanol (80% V/V), using sinigrin (2-propenylglucosinolate) as internal standard. The extract was loaded onto Sephadex DEAE A25 columns, previously equilibrated in 0.02 M acetate buffer (pH 5), and incubated overnight with *Helix pomatia* Type H-1 sulfatase (S9626, Sigma-Aldrich) prepared as described by Graser et al (Graser et al., 2000). Desulfoglucosinolates were eluted in H2O and separated by HPLC (Agilent HPLC 1200 series) on a C-18 reversed phase column (Supelcosil LC-18, L × i.d. 250 × 2.1 mm; particle size 3 μm) at 25°C using a H2O-acetonitrile (solvent A-solvent B) gradient (0—2 min 3% B; 2—17 min 3-40% B, 17-22 min 40% B, 22—22.1 min 40—100% B, 22.1—32 min 100% B, 32—32.1 100-3% B, 32.1—60 min 3% B) at a flow rate of 0.3 ml min^-1^. Desulfoglucosinolates were monitored by UV diode array detection, identified based on retention times and LC-MS confirmation of peaks, quantified by A229nm relative to the standard and taking into account the response factors (Brown et al., 2003).

### Statistical analysis

Collected data from different experiments were analyzed using Microsoft Excel. ANOVA and mean comparisons were performed using Agricolae package in R environment (De Mendiburu, 2014). qRT-PCR data were analyzed using the qBase^+^ software (Version 3.2, Biogazelle).

## Results

### PIPL6 is highly inducible by different classes of elicitors

To draw a comprehensive picture of the dynamics of *PIPL6* expression to plant immune response elicitors, a set of time course experiments were executed. Wild-type Arabidopsis seedlings were treated with two well-known PAMPs, flg22 peptide (100 nM) and chitin hexamers (500 nM), and PEP1 peptide (100 nM) as a DAMP elicitor. Expression analysis using qRT-PCR showed that the expression of *prePIPL6* starts to increase 15 min after treatment and culminated between 30-60 min (Fig. 1). Among the different classes of elicitors, the flg22 treatment exhibited the strongest effect on *prePIPL6* expression while chitin and PEP1 showed similar effects, with a more durable induction in response to the PEP1 treatment.

**Figure 1.**
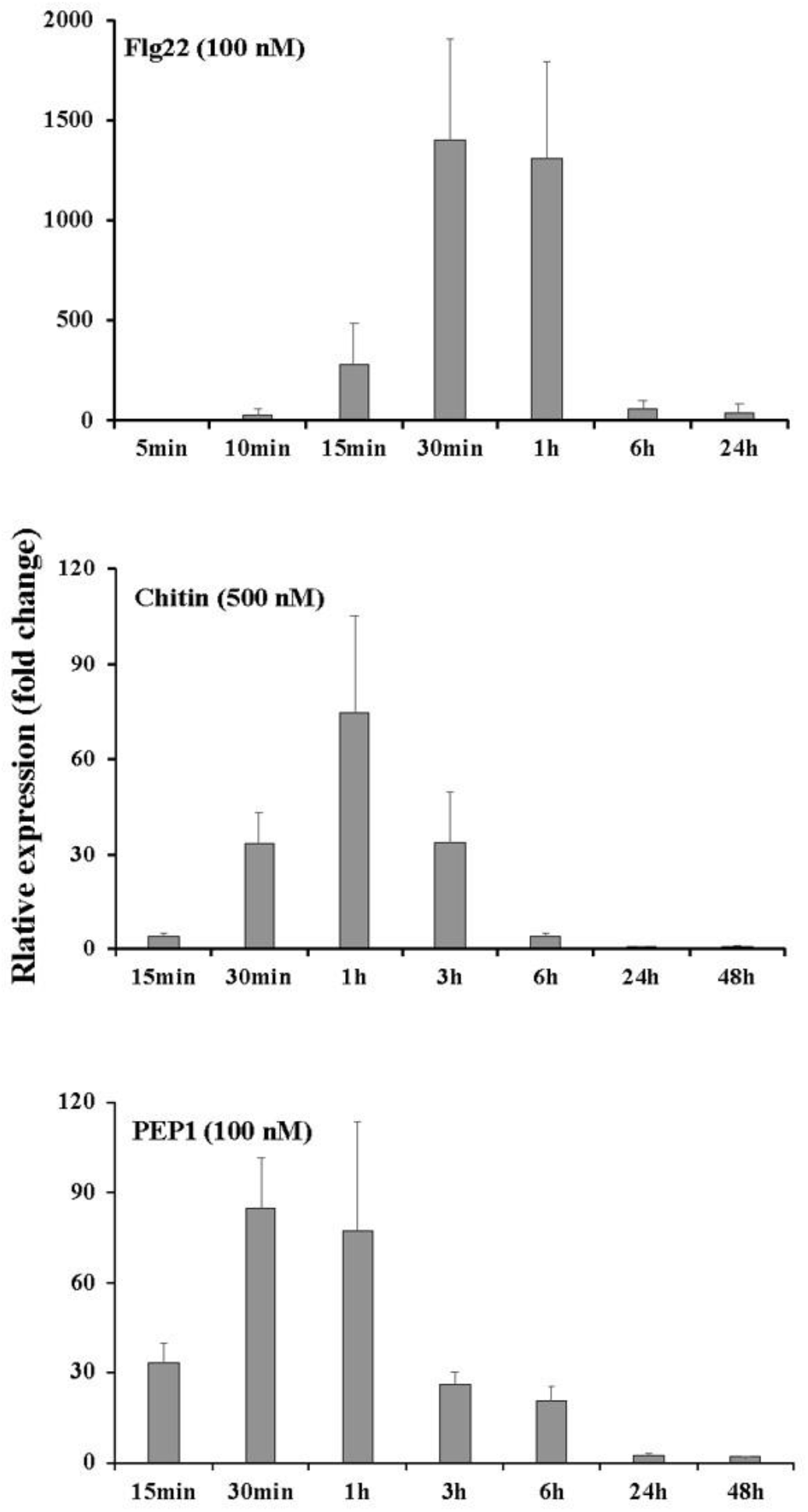
*PIPL6* expression is highly and transiently inducible in response to exogenous application of plant immune elicitors. Two-weeks-old seedlings were treated by different class of elicitors or water as control and tissues were harvested at different time points for RNA isolation and qRT-PCR analyses. Expression levels were calculated for each time point relative to the control treatment. Bars and error bars represent mean and standard deviations, calculated from three biological replicates, respectively.

### Ectopic application of PIPL6 synthetic peptide causes massive transcriptome reprogramming

In order to gain an insight about the downstream biological processes and pathways affected by the PIPL6 peptide, we designed a synthetic peptide containing the C-terminal conserved domain of PIPL6 (last 14 amino acids) including the conserved SGPS motif. Two-weeks-old Wt plants were treated with 1 μM synthetic peptide for three hours, and rosette leaves were harvested for global transcriptome sequencing. Interestingly, 753 genes were found differentially regulated (587 genes upregulated and 166 downregulated) using stringent statistical thresholds (2-fold or more compared with control treatment, *P*≤0.01). Gene Ontology enrichment of the top 100 upregulated genes revealed that the synthetic peptide treatment affected biological pathways involved in detoxification processes, plant responses to biotic and abiotic stresses, metabolite homeostasis, and production of indolic glucosinolates (Fig. 2A). A detailed survey on the upregulated genes revealed many known immune marker genes, including the *FRK1, RLK7*, and *NILR1* receptors, *PAD3* and *CYP71A12* in the camalexin biosynthesis pathway, *CYP81F12*, and *MYB51* in the glucosinolate biosynthesis pathway, and many TFs including *WRKY18, WRKY40, WRKY60, WRKY33, WRKY53* (Table 1 and Supplementary Table 2). To confirm the biological relevance of the PIPL6 effect on plant immune response pathways, the peptide treatment was repeated under the same conditions and RNA was subjected to qRT-PCR analysis of some selected marker genes from different pathways (Fig. 2B). Induction of *RLK7* expression upon PIPL6 treatment prompted us to examine if RLK7 functions as a receptor in PIPL6 initiated gene expression regulation. To test this hypothesis, Wt and *rlk7* seedlings were treated with the PIPL6 peptide and the expression of marker genes were studied using qRT-PCR. Induction of the selected genes in response to PIPL6 in Wt seedlings were found to be suppressed in the *rlk7* background (Fig. 2C). This observation suggests that RLK7 plays an important role as a receptor for PIPL6 initiated signalling.

**Figure 2.**
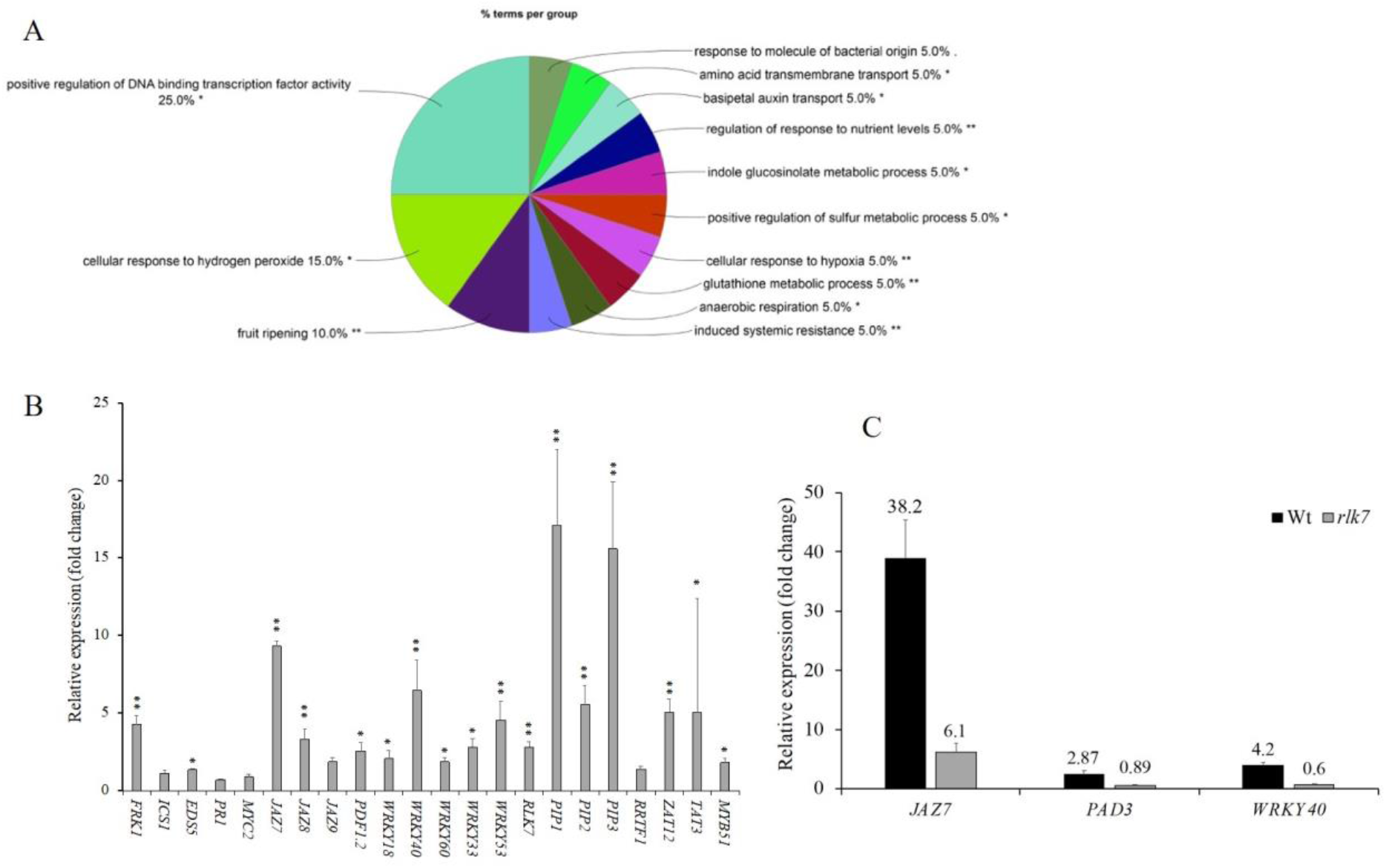
Exogenous application of PIPL6 synthetic peptide elicits plant immune response in an RLK7 dependent manner. A) GO enrichment of top 100 upregulated genes 3h after PIPL6 treatment of Wt plants followed by RNA-seq analysis. B) Verification of RNA-seq data by qRT-PCR. The whole treatment was repeated and tissue were used for RNA isolation and qRT-PCR analyses for the selected genes. Bars and error bars represent means and standard deviations calculated from four biological replicates. * and ** represent statistically significant differences compared to the control at *p-value* < 0.05 and <0.01 levels respectively. C) Induction of plant immune response by PIPL6 synthetic peptide requires RLK7. Wt and *rlk7* mutant plants were treated by PIPL6 peptides and tissues were harvested for RNA isolation and qRT-PCR for the selected marker genes. Bars and error bars represent means and standard deviations calculated from four biological replicates. Mean expression values indicated on top of each bar.

**Table 1.** List of differentially regulated genes and pathways three hours after synthetic PIPL6 treatment.

| Gene ID | Gene name and Pathway | Log2FC | P-value |
| --- | --- | --- | --- |
| **Detoxification and redox homeostasis** |  |  |  |
| AT2G04050.1 | AtDTX3 (Protein DETOXIFICATION 3) | 6,057 | 8,1E-112 |
| AT2G04040.1 | AtDTX1 (MATE efflux family protein) | 4,666 | 5,4E-84 |
| AT5G62480.1 | ATGSTU9 (glutathione S-transferase tau 9) | 5,528 | 2,3E-74 |
| AT2G04070.1 | AtDTX4 (Protein DETOXIFICATION 4) | 5,176 | 4,9E-58 |
| AT2G29470.1 | ATGSTU3 (glutathione S-transferase tau 3) | 4,986 | 7,8E-06 |
| AT2G29460.1 | ATGSTU4 (glutathione S-transferase tau 4) | 3,743 | 7,9E-29 |
| AT1G02930.1 | ATGST1 (glutathione S-transferase 1) | 3,342 | 3,3E-78 |
| AT5G02780.1 | GSTL1 (glutathione transferase lambda 1) | 3,049 | 9,4E-06 |
| AT1G02920.1 | ATGST11 (glutathione S-transferase 11) | 2,864 | 1,5E-52 |
| AT1G17170.1 | ATGSTU24 (glutathione S-transferase tau 24) | 2,362 | 4,9E-62 |
| AT2G02930.1 | ATGSTF3 (glutathione S-transferase F3) | 2,211 | 1,2E-06 |
| **Glucosinolates biosynthesis** |  |  |  |
| AT5G57220.1 | CYP81F2 | 2,663 | 1,0E-76 |
| AT2G30750.1 | CYP71A12 | 2,511 | 5,4E-19 |
| AT1G18570.1 | AtMYB51 | 1,322 | 2,5E-24 |
| **Camalexin** |  |  |  |
| AT3G26830.1 | PAD3 (PHYTOALEXIN DEFICIENT 3) | 4,501 | 1,5E-47 |
| AT2G30750.1 | CYP71A12 | 2,511 | 5,4E-19 |
| **Immunity related receptors** |  |  |  |
| AT1G09970.2 | RLK7 (receptor-like kinase 7) | 1,395 | 1,2E-35 |
| AT2G19190.1 | FRK1 (FLG22-induced receptor-like kinase 1) | 1,832 | 1,0E-20 |
| AT1G74360.1 | NILR1/GRACE | 1,946 | 2,0E-41 |
| AT2G39660.1 | BIK1 (botrytis-induced kinase1) | 0,897 | 6,4E-17 |
| AT5G25910.1 | AtRLP52 (receptor like protein 52) | 2,183 | 3,1E-10 |
| AT2G32680.1 | AtRLP23 (receptor like protein 23) | 1,876 | 2,8E-05 |
| AT3G05360.1 | AtRLP30 (receptor like protein 30) | 0,994 | 3,4E-08 |
| **JA** |  |  |  |
| AT2G34600.1 | JAZ7 (jasmonate-zim-domain protein 7) | 3,212 | 4,2E-09 |
| AT1G19180.2 | JAZ1 (jasmonate-zim-domain protein 1) | 1,419 | 1,4E-12 |
| AT3G25780.1 | AOC3 (allene oxide cyclase 3) | 1,058 | 7,1E-07 |
| **Ethylene** |  |  |  |
| AT4G26200.1 | ACS7 (1-amino-cyclopropane-1-carboxylate synthase 7) | 2,766 | 7,1E-16 |
| AT1G01480.1 | ACS2 (1-amino-cyclopropane-1-carboxylate synthase 2) | 3,072 | 2,9E-12 |
| AT4G08040.1 | ACS11 (1-aminocyclopropane-1-carboxylate synthase 11) | 2,164 | 1,4E-15 |
| AT4G11280.1 | ACS6 (1-aminocyclopropane-1-carboxylic acid synthase 6) | 1,298 | 9,5E-18 |
| AT2G47520.1 | ERF71; HRE2 (HYPOXIA RESPONSIVE ERF | 4,015 | 5,3E-30 |
| AT1G43160.1 | ERF113 ; RAP2.6 (related to AP2 6) | 3,446 | 4,0E-11 |
| **WRKY TFs** |  |  |  |
| AT1G80840.1 | WRKY40 | 2,976 | 4,3E-37 |
| AT5G13080.1 | WRKY75 | 2,562 | 1,5E-05 |
| AT4G23810.1 | WRKY53 | 1,846 | 1,4E-36 |
| AT2G38470.1 | WRKY33 | 1,505 | 3,4E-42 |
| AT2G25000.1 | WRKY60 | 1,328 | 4,6E-09 |
| AT5G49520.1 | WRKY48 | 1,288 | 2,3E-05 |

### PIPL6 synthetic peptide activates the MAPK6, MAPK3 and WRKY33 module in an RLK7 dependent manner

Mitogen-activated protein kinase (MAPK) cascades are recognized as one of the most conserved signal transduction modules in eukaryotes that function as nodes of convergence and divergence for many endogenous and exogenous triggered signal transduction pathways (Ligterink et al., 1997). To investigate if PIPL6 can activate a MAPK cascade, Wt and *rlk7* seedlings were treated with synthetic PIPL6 and flg22 peptides. Immunoblotting using a phosphoantibody against MAPK6, MAPK4, and MAPK3 revealed that unlike flg22, which activates all three MAPKs, PIPL6 peptide only activates MAPK6 and MAPK3. Our assay showed that both MAPK6 and MAPK3 are activated by phosphorylation 5 minutes after PIPL6 treatment, culminating after 15 minutes. The signal was attenuated after 60 minutes. Furthermore, PIPL6 peptide activation of MAPK6 and MAPK3 was abolished in the *rlk7* background, supporting the function of RLK7 as a receptor for PIPL6 (Fig. 3a). Using a similar experimental set-up, we applied synthetic flg22 and PIPL6 peptides to the *wrky33* mutant and transgenic plants expressing a WRKY33:HA construct transformed in *wrky33* mutant background to investigate their effects on the induction of WRKY33. Immunoblotting using an anti-HA antibody showed that both flg22 and PIPL6 peptides are able to activate WRKY33 to the same extent (Fig. 3b).

**Figure 3.**
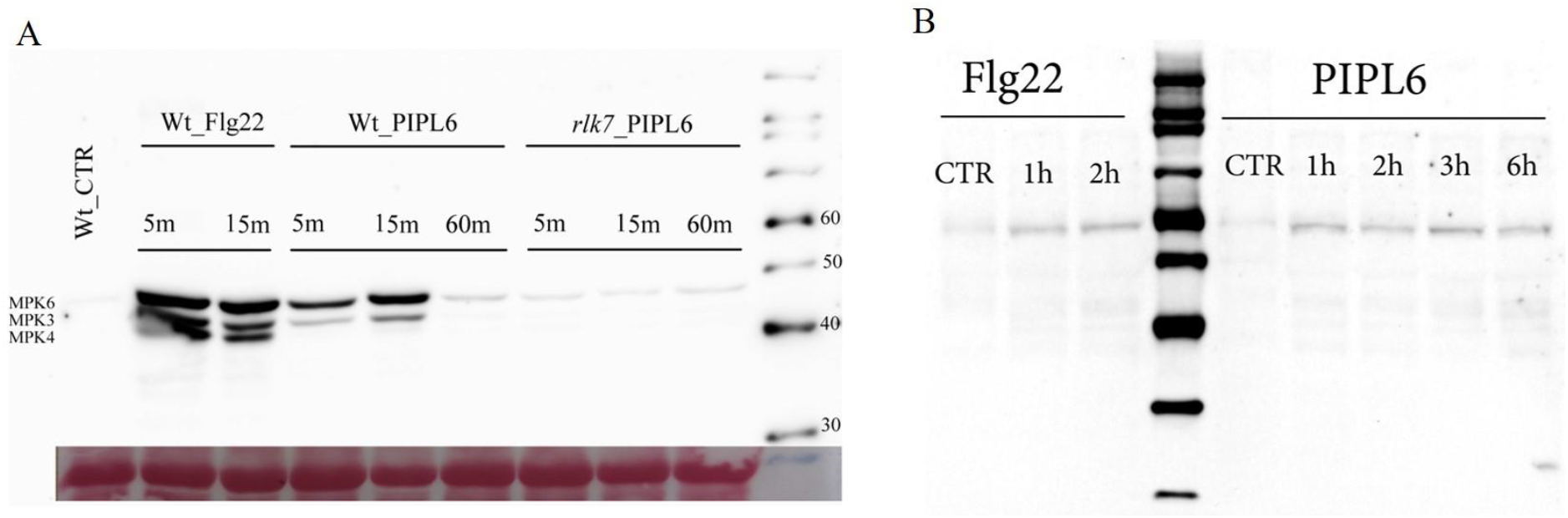
Exogenous application of PIPL6 synthetic peptide activates MPK6/3 and WRKY33 module. A) Phosphorylation of MAP6/3 in response to flg22 and PIPL6 elicitors in Wt and *rlk7* mutant seedlings. Seedlings of Col-0 (Wt) and *rlk7* mutants were treated with water for 5min as control or with 1 μMol flg22 or PIPL6 for the indicated times. Protein samples were subjected to immunoblot analysis using antibodies against phospho-p44/p42 (α-pTEpY), MPK3, and MPK6. Lower panel is the Ponceau Red staining of the same blot that used as loading control. The experiment was repeated three times with similar results. B) Trasngenic seeds expressing *proWRKY33*:WRKY33:HA construct in *wrky33* background were grown and treated by flg22 and PIPL6 synthetic peptides at different time points as explained above. Proteins were isolated from samples and subjected to immunoblot analysis with Anti-HA antibody.

### WRKY33 is the dominant transcription factor regulating PIPL6 expression

Previously, we showed that a network of WRKY TFs including WRKY18, WRKY40, WRKY60 and WRKY33 actively regulate expression of *PIP1-3* genes during flg22 treatment and *Botrytis cinerea* infection (Najafi et al., 2020). Sequence analysis of the *PIPL6* promoter region revealed the presence of two W-boxes at −155 and −744 positions upstream of the start codon, which might act as binding sites for WRKY TFs (Fig. 4a). To further analyze the effects of the above mentioned TFs, we analysed the expression levels of *PIPL6* in *wrky18, wrky40*, and *wrky60* single and double knock-outs, as well as *wrky33* mutant. No significant effects in the expression levels of *PIPL6* were found in *wrky18*, *40, 60* single knock-out mutants (Fig. 4b). In contrast, all three combinations of double knock-out mutants for WRKY18, WRKY40 and WRKY60 exhibited significant effects as positive regulators of *PIPL6* expression. However, *PIPL6* expression was reduced about 85% in the *wrky33* mutant this points at WRKY33 as a positive regulator.

**Figure 4.**
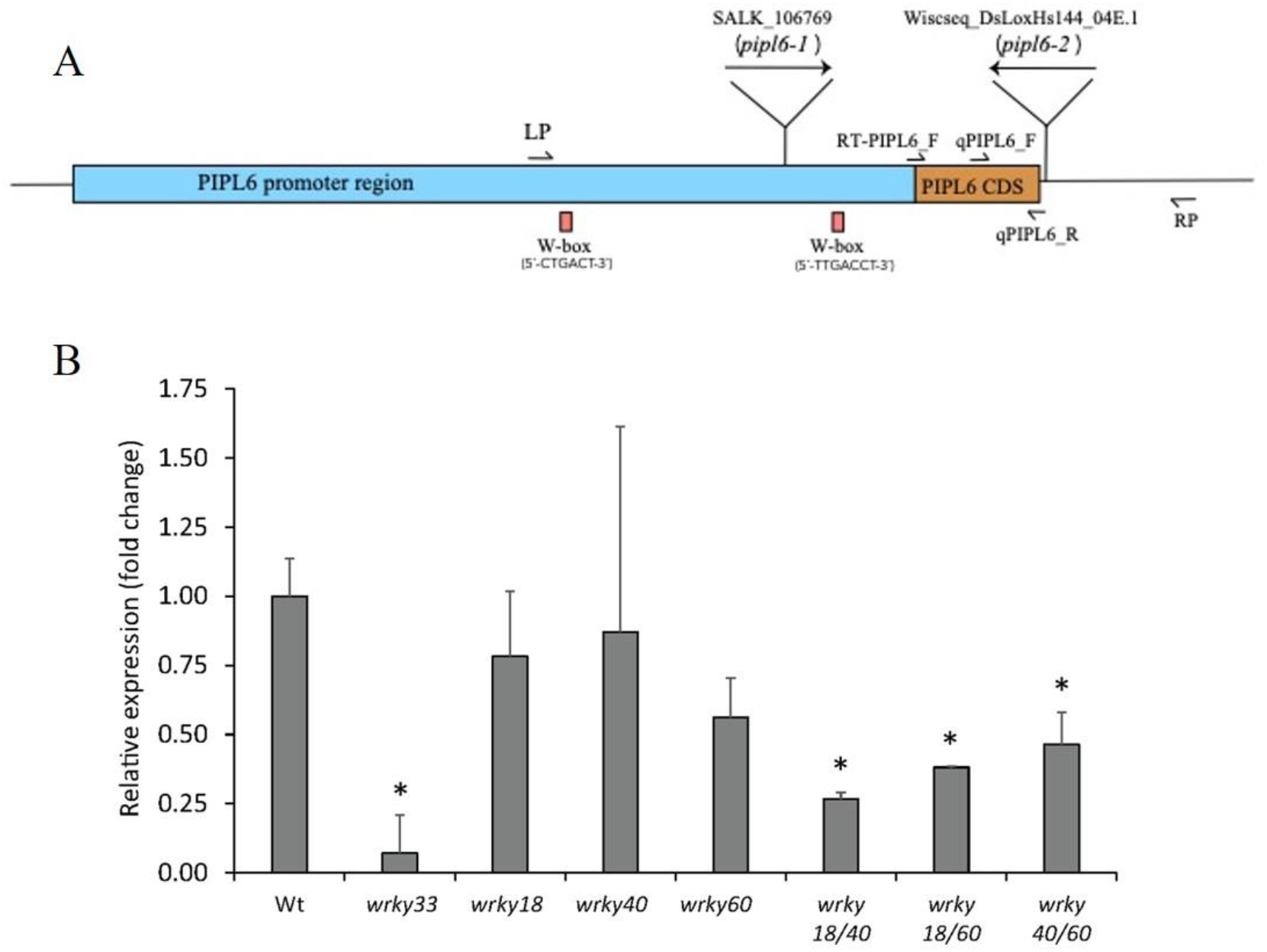
A) Genomic map, T-DNA insertion sites and the position of W-boxes in the *PIPL6* promoter region relative to the start codon. LP and RP primers were used for screening of T-DNA lines. RT-PIPL6_F in combination of qPIPL6_R primers were used to screen the mutants at mRNA levels. qPIPL6_F and qPIPL6_R primers were used in qRT-PCR analyses. B) Combination of WRKY TFs function as positive regulator of *PIPL6* expression. Wt, different single knock-outs, and combination of double knock-out seeds were grown for seven days and treated by 1μM of flg22 or water as control and tissue harvested 1h after treatment. Relative expressions were normalized against the Wt response. Bars and error bars represent mean and standard deviations calculated from three biological replicates. * indicate a statistically significant differences at *p-value*<0.05.

### Characterization of mutant and overexpression lines

The PIPL6 encoding gene is not annotated on Arabidopsis genome and hence there is no information about its function *in-silico*. The PIPL6 encoding sequence is located on the chromosome 1 and we have reported a tentative gene ID (At1g47178) for it (Vie et al., 2015). To investigate the biological function of the PIPL6 peptide, we mapped all available T-DNA and transposon insertion lines around *PIPL6* encoding genomic region. Two independent putative knock-down lines were identified. The SALK_106769 line, which has T-DNA inserted in the promoter region at the position of −180 upstream of start codon, was designated as *pipl6-1*. The Wisc_DsLoxHs144_04E.1 line, with a transposon inserted immediately after the stop codon, will be referred to as *pipl6-2* (Fig. 4a). Homozygosity of selected lines was confirmed at both genomic and transcription levels. Due to very low levels of *PIPL6* expression under normal conditions, we treated Wt and *PIPL6* knock-down lines using flg22, and tissue was harvested 1h after treatment for RNA isolation and RT-PCR. As expected, some degree of expression was detected before and after flg22 treatment for the *pip6-1* line.

Transgenic plants overexpressing *PIPL6* coding sequence under control of the constitutive CaMV35S promoter were generated. Two independent T3 lines with constitutive expression of *PIPL6* were chosen for further analysis. Under normal growth conditions, no visible growth or developmental abnormalities were observed in any of the lines (Supp Fig. 1). Overexpression of *PIPL6* was confirmed using RT-PCR in selected lines.

### Manipulation of *PIPL6* expression alters plant responses to necrotrophic fungal pathogens

Leaves of five-weeks-old Wt, *wrky33, pipl6* knock-down, and *PIPL6:OX* lines were inoculated with a *Botrytis cinerea* isolate 2100 (Spanish type) spore suspension. Wt Col-0 and *wrky33* mutant plants have been reported to be resistant and susceptible to this strain, respectively (Birkenbihl et al., 2012). In Wt and *PIPL6:OX* lines, inoculation caused lesions at the site of infection two days after treatment, but development of necrotic lesions halted after day 3 (Fig. 5a.). In contrast, the symptoms developed much faster in *wrky33* and *pipl6* knock-down plants. Further observations of lesion development revealed that background symptoms developed faster in *wrky33* than in *pipl6* knock-down lines and caused complete destruction of infected plants 7 dpi. In *pipl6* knock-down lines, the symptom development rate and severity were lower than in *wrky33* plants. Measurements of lesion size in inoculated detached leaves four days after inoculation revealed that *pipl6* knock-down plants displayed larger necrotic lesions compared to the Wt plants (Fig. 5b.). However, this *Botrytis cinerea* strain is not able to discriminate the potential resistant phenotype caused by the overexpression of *PIPL6*. To investigate this hypothesis, we challenged all genotypes by spraying spores from *Alternaria brassicae* and assessing their responses five dpi. Similar to Botrytis, three dpi, all genotypes exhibited small lesions at the sites of spore contact. However, compared to the Wt plants, lesions developed faster in *wrky33* and more slowly in *pipl6* knock-down lines. In both *PIPL6:OX* lines, the development of symptoms was halted three dpi and plants showed a resistant phenotype (Fig. 6 and Supplementary Fig. 2).

**Figure 5.**
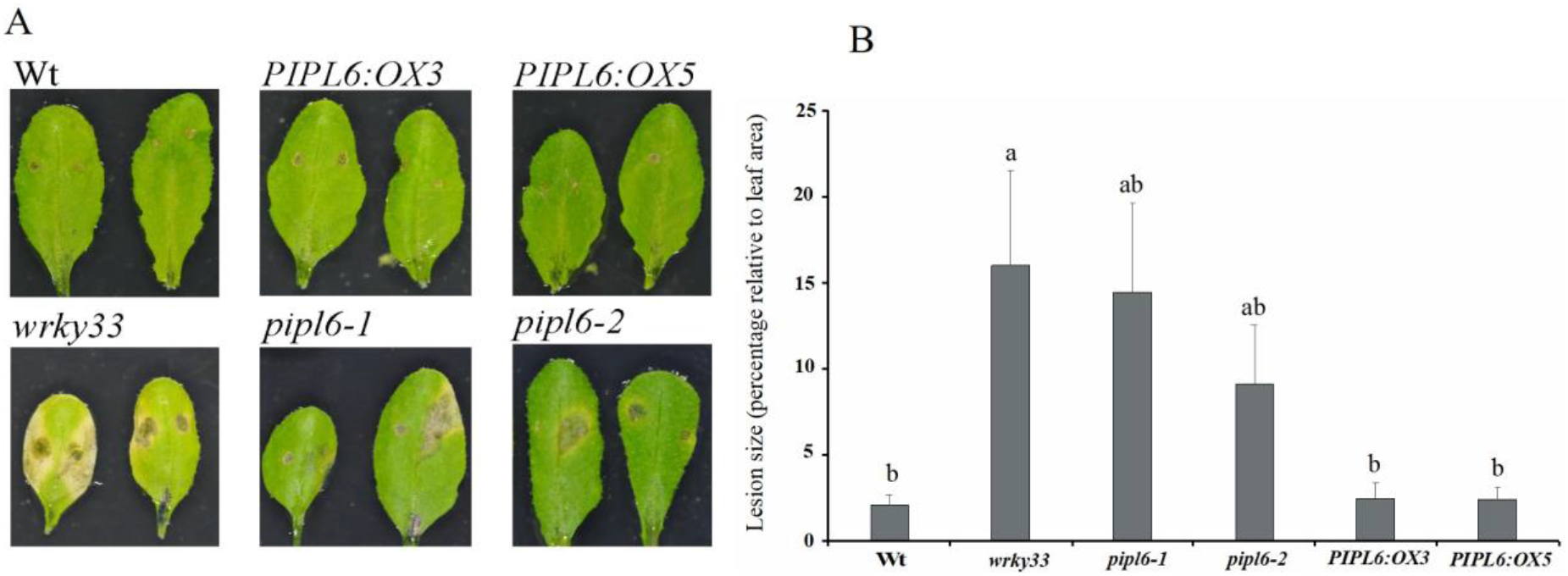
Down-regulation of PIPL6 increase plant susceptibility towards *Botrytis cinerea* infection. A) Symptoms development of rosette leaves inoculated by *Botrytis cinerea* spores three days post infection. B) Measurement of lesion sizes relative to the whole leaf area four days post infection. Bars and error bars represent means and standard deviations calculated for inoculated leaves for each genotype. Different letters on bars indicate statistically significant differences calculated by ANOVA followed by Tukey multiple mean comparison method at *p-value* > 0.05, n>10.

**Figure 6.**
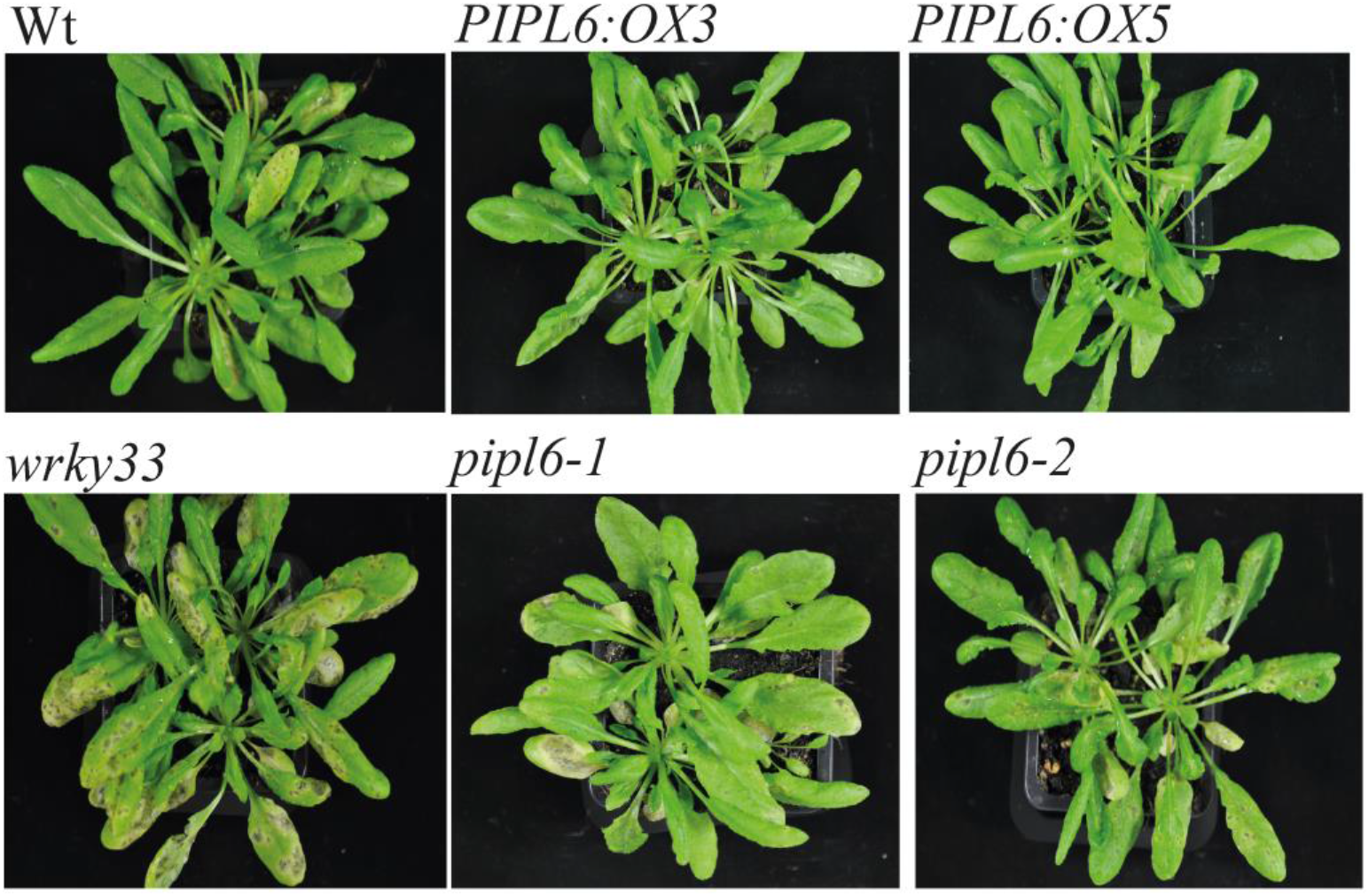
Overexpression of *PIPL6* enhances plant resistance against *Alternaria brassicae* infection. Five-weeks-old Wt and transgenic lines were sprayed by *Alternaria brassicae* spores and plants were photographed five days after inoculation.

### Expression of key genes in JA signalling and camalexin biosynthesis are differentially regulated in *PIPL6* transgenic lines during Botrytis infection

Previous studies have shown that infection of Arabidopsis by Botrytis causes a massive transcriptional reprogramming of many genes including those that are involved in the biosynthesis and signalling of JA, SA, ethylene, ABA, and camalexin (Birkenbihl et al., 2012). The altered responses of *PIPL6* transgenic lines to Botrytis and the differential regulation of JA and camalexin marker genes after applying synthetic PIPL6 peptide prompted us to investigate the expression of key genes in these pathways upon Botrytis infection in *PIPL6* lines. *wrky33* mutant and Wt plants were included in the assays as susceptible and resistant lines respectively. Five-week-old short days grown plants were inoculated by Botrytis spores and rosette leaves were sampled 48 hpi for gene expression and metabolite analysis. WRKY33 is the main TF regulating *PIPL6* expression (Fig. 4b.). On the other hand, ectopic application of PIPL6 synthetic peptide induces *WRKY33* expression (Fig. 2b. and Table 1). Analysis of *WRKY33* expression in *PIPL6* transgenic lines after Botrytis infection revealed that expression of *WRKY33* is significantly higher in *PIPL6:OX* lines compared to *pipl6* mutants and Wt plants (Fig. 7a.). Expression of *ICS1*, the key gene in SA biosynthesis, and *PR1*, a marker gene for the SA pathway, was highest in the *wrky33* background. *ICS1* expression did not differ significantly in *pipl6* mutants and *PIPL6:OX* overexpression lines compared to the Wt plants (Fig. 7b.). However, *PR1* expression was significantly higher in *PIPL6:OX* lines than in *pipl6* mutants and Wt plants (Fig. 7a.).

**Figure 7.**
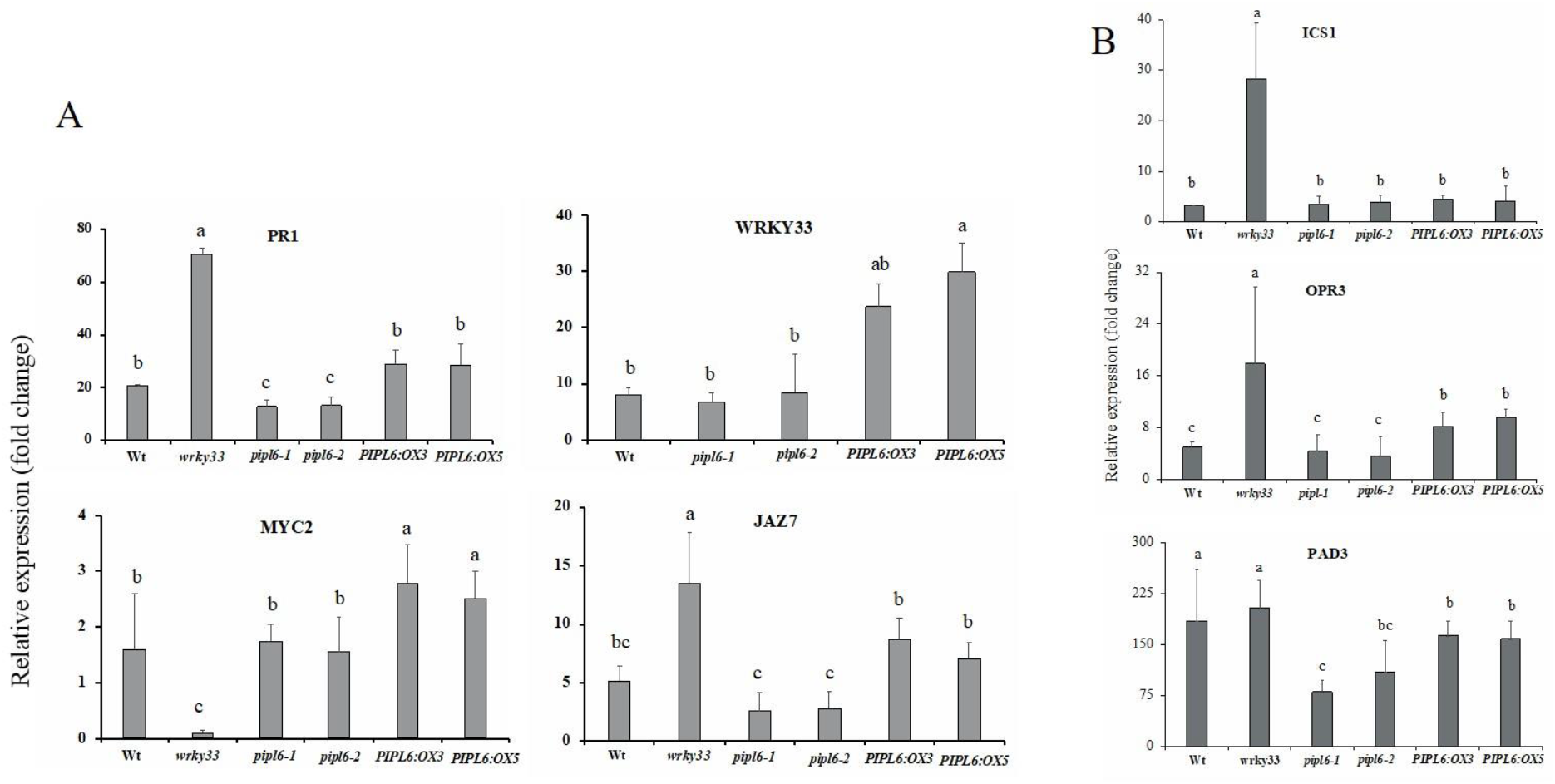
Expression of key genes involved in SA, JA, and camalexin production and signaling pathways are differentially regulated in *PIPL6* transgenic lines upon *Botrytis cinerea* infection. Five-weeks-old plants were challenged by Botrytis spores and rosette leaves were harvested 48hpi for RNA isolation and qRT-PCR analysis. Bars and error bars represent means and standard deviations calculated from four biological replicates for each genotype. Different letters on bars indicate statistically significant differences calculated by ANOVA followed by Tukey multiple mean comparison method at *p-value* > 0.05.

Gene expression was analyzed for *OPR3* as a marker for JA biosynthesis and for *MYC2* and *JAZ7* as markers of JA signaling. *OPR3* expression was significantly higher in the *wrky33* background followed by *PIPL6:OX* lines, but there were no significant differences between *pipl6* mutants and Wt lines (Fig. 7b.). Expression of *MYC2*, the major TF of JA responsive genes, was significantly higher in *PIPL6:OX* lines than in Wt and *pipl6* mutant lines, while it was repressed in the *wrky33* background (Fig. 7a.). *JAZ7* expression pattern exhibited differential response in *PIPL6:OX* and mutant lines: slightly upregulated in overexpression plants and downregulated in mutant lines (Fig. 7a.). The expression pattern of *PAD3* showed a differential regulation upon Botrytis infection in *PIPL6:OX* and *pipl6* knock-down lines. *PAD3* expression was significantly low in overexpression lines compared to Wt and *wrky33* plants while its expression was induced even in lower extend in mutant lines (Fig. 7b).

### Overexpression of *PIPL6* is associated with induction of jasmonic acid and loss of function of PIPL6 represses camalexin biosynthesis during Botrytis infection

SA, JA/ET and ABA production, signaling and their cross-talk determine plant defense responses against necrotrophic pathogens including *Botrytis cinerea* and *Alternaria brassicae* (Berens et al., 2017). In addition to phytohormones, production of camalexin is also a key player in mounting the proper immune response in Arabidopsis in response to pathogens. We therefore measured SA, JA, ABA, and camalexin levels in Wt, *wrky33*, and *PIPL6* transgenic lines upon Botrytis infection. Under mock treatment, no significant deviations in production of SA, JA, ABA, and camalexin were observed between *PIPL6* overexpression and mutant lines compared to the Wt. However, in the *wrky33* background SA, JA and camalexin levels were higher than in all other tested lines (Supplementary Fig. 3). When plants were challenged by Botrytis spores, SA levels increased in all lines at 48 hpi, particularly in the *wrky33* background. SA levels in *PIPL6:OX* lines were induced to a similar extent than in Wt plants while the SA level was less induced in *pipl6-1* plants (Fig. 8). JA production was also induced at 48 hpi, but displayed a differential regulation in PIPL6 transgenic lines. While *PIPL6:OX* plants produced significantly higher levels of JA than Wt, *pipl6-1* plants produced lower levels than Wt when challenged by Botrytis spores (Fig. 8). Previous studies have shown that *wrky33* susceptibility is negatively associated with ABA production upon Botrytis infection (Liu et al., 2015). To investigate the susceptibility observed in *pipl6-1* mutant plants we measured the ABA production in infected plants. As expected, ABA production in *wrky33* plants was significantly induced, but none of the *PIPL6* overexpression and mutant plants showed significant inductions increase in ABA levels compared to the Wt plants (Fig. 8). Furthermore, we quantified camalexin levels in the studied genotypes. During Botrytis infection WRKY33 directly binds to the promoter region of *PAD3* and regulates camalexin production. Hence, the amount of camalexin in *wrky33* plants remained significantly lower than in all other lines. We found that downregulation of *PIPL6* also led to a significantly lower induction in camalexin production, while *PIPL6* overexpression did not alter camalexin levels when compared to the Wt plants (Fig. 8).

**Figure 8.**
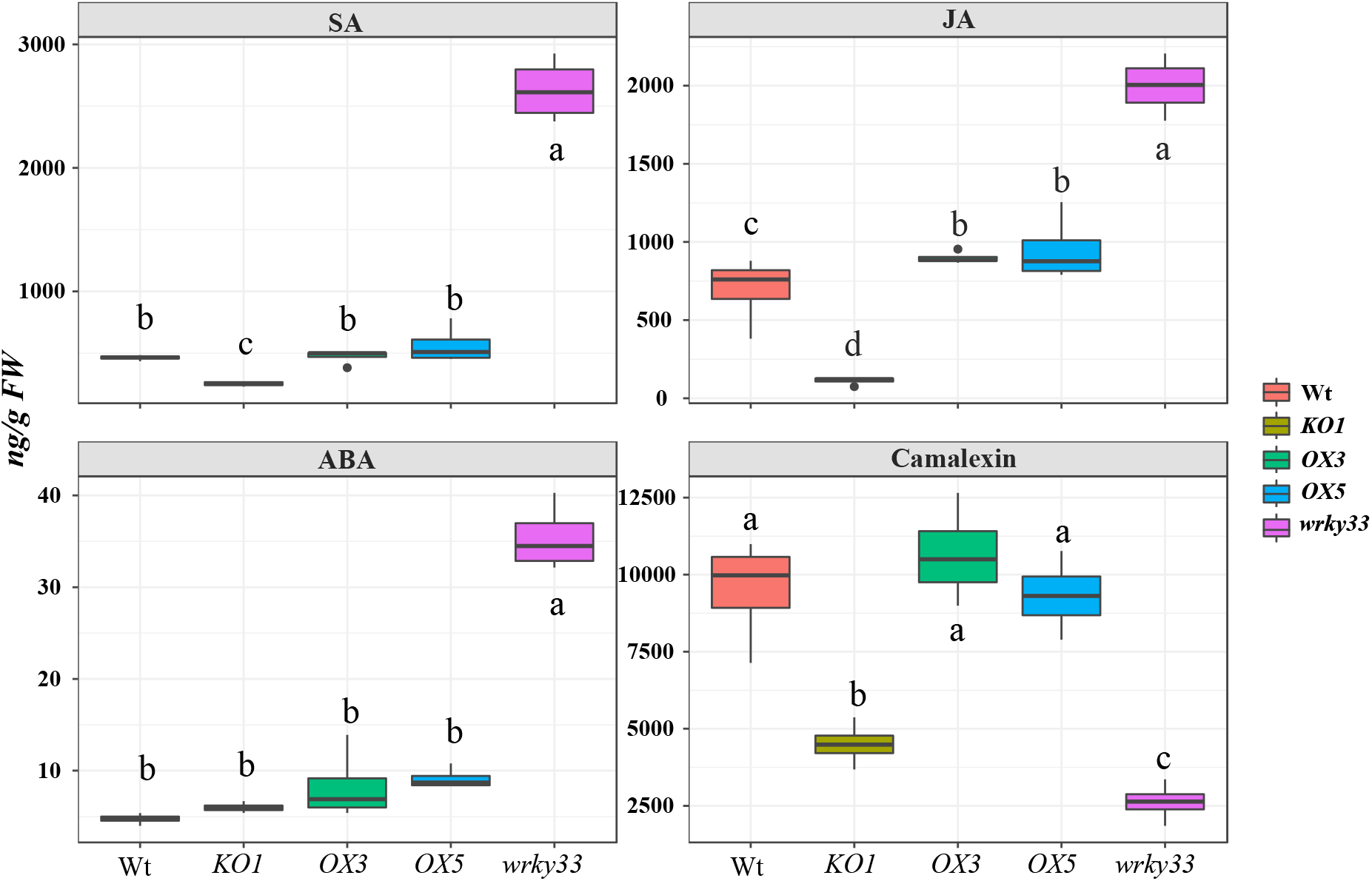
Productions of SA, JA, and camalexin are impaired in *pipl6* knock-down line in response to *Botrytis cinerea* infection. Five-weeks-old plants were challenged by Botrytis spores and rosette leaves were harvested 48hpi for hormonal extraction and quantification. Different letters indicate statistically significant differences calculated by ANOVA followed by Tukey multiple mean comparison method (n = 4, *p-value* < 0.05).

### *PIPL6* expression affects glucosinolate profiles upon Botrytis infection

Glucosinolates are sulfur-containing secondary metabolites that are present in Brassicaceae (Blažević et al., 2020). We therefore monitored the expression of MYB28 and MYB51, two TFs that are important for regulating the biosynthesis of aliphatic and indolic glucosinolates respectively (Gigolashvili et al., 2007), 48 h after Botrytis Infection. Repression of *MYB28* expression by Botrytis infection was similar in Wt and *PIPL6:OX* backgrounds, but was significantly stronger in *wrky33* plants. Interestingly, the least repression of *MYB28* expression was observed in *pipl6-1*, while in *pipl6-2* mutants the repression was higher that Wt plants. *MYB51* expression, on the other hand, showed a significant induction in *wrky33* and *PIPL6:OX* lines compared to the *pipl6* mutants and Wt (Fig. 9a.). These changes in *MYB28* and *MYB51* expression patterns prompted us to measure glucosinolate levels at this time point. Total glucosinolate levels were lower in Wt plants 48 hpi when compared to the mock-treated plants. *wrky33* and the two *pipl6* mutants showed even lower total amounts of glucosinolates than Wt when exposed to Botrytis. However, PIPL6:OX3 plants produced similar levels as Wt. Reduced levels of both aliphatic and indolic glucosinolate contributed to lower total amounts of glucosinolates in the mutant lines (Fig. 9b.). Differences between *PIPL6:OX* lines were however due to changes in aliphatic glucosinolates, while levels of indolic glucosinolates were similar among them. As 4-methylsulfinylbutylglucosinolate (4MSB) was the most abundant aliphatic glucosinolate, changes in its levels had the biggest impact, but similar patterns were observed for the other aliphatic glucosinolates, except 4-methylthiobutylglucosinolate (4MTB). The three indolic glucosinolates showed more varied patterns. While all *PIPL6* lines showed similar levels of indol-3-ylmethylglucosinolate (I3M) and 4-methoxyindol-3-ylmethylglucosinolate (4MeO-I3M) than Wt when exposed to Botrytis, the levels of 1-methoxyindol-3-ylmethylglucosinolate (1MeO-I3M) were significantly different from those in Wt. No substantial differences were observed for individual indolic glucosinolates between the *PIPL6* lines (Supplementary Fig. 4).

**Figure 9.**
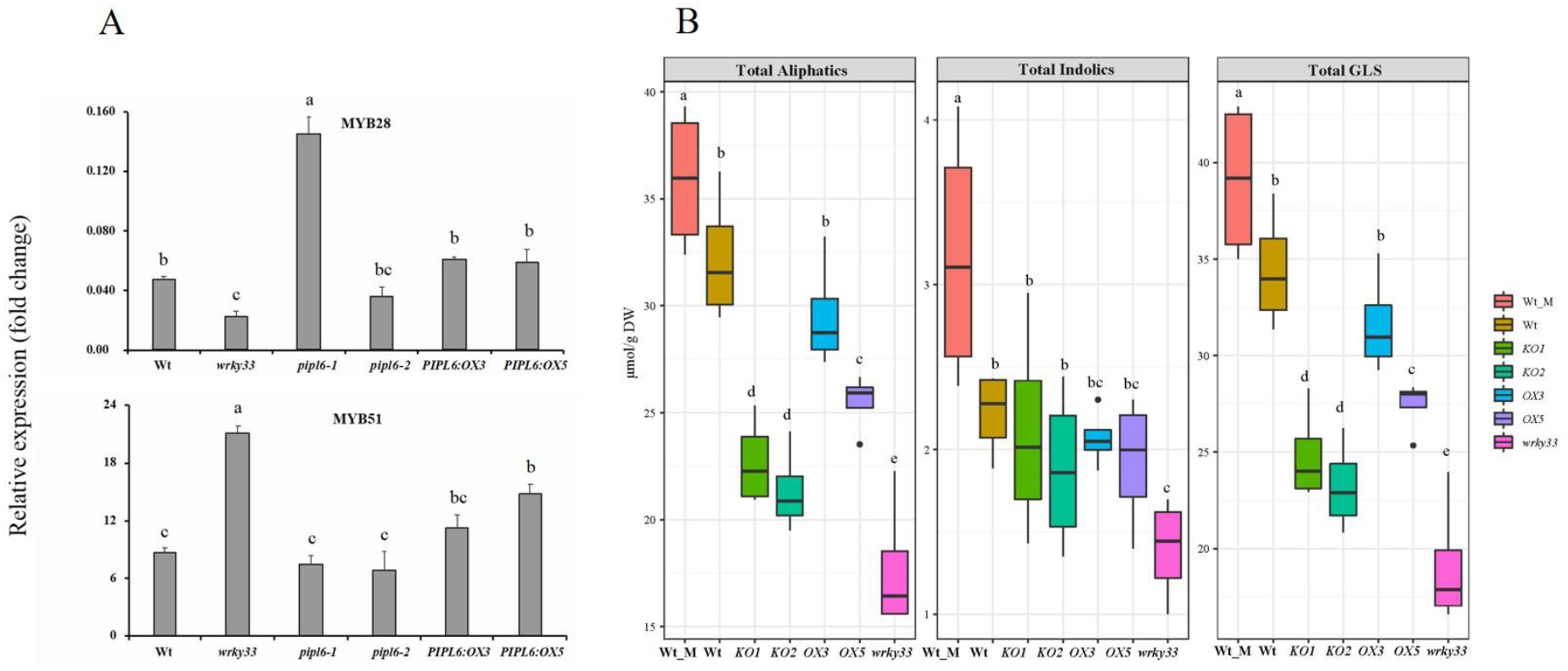
Alteration in *PIPL6* expression affects glucosinolates production pathways. A) Expression pattern of two key TFs involved in glucosinolates production in different genotypes 48h after *Botrytis cinerea* infection. Bars and error bars represent means and standard deviations calculated from four biological replicates for each genotype. Different letters on bars indicate statistically significant differences calculated by ANOVA followed by Tukey multiple mean comparison method at *p-value* > 0.05. B) Glucosinolates production profiles in different genotypes 48h after *Botrytis cinerea* infection. Different letters indicate statistically significant differences calculated by ANOVA followed by Tukey multiple mean comparison method (n = 4, *p-value* < 0.05).

## Discussion

Among several classes of endogenous elicitors, signalling peptides have been investigated for their role in regulating plant immunity (Hu et al., 2018). We have reported a new class of peptides termed as PAMP-INDUCED SECRETED PEPTIDE (PIP) and PIP-LIKE (PIPL), which shows similarity to the IDA/IDA-LIKE and C-TERMINALLY ENCODED PEPTIDE (CEP) peptides (Vie et al., 2015). It has been shown that PIP1 functions as an amplifier of plant immune response against biotrophic and necrotrophic pathogens through RLK7 as the main receptor (Hou et al., 2014). A recent study showed that an ortholog of PIP1 in potato (*StPIP1*) positively regulates the immune response to *Potato virus Y* (PVY) infection (Combest et al., 2021). Among the eleven members in the PIP/PIP-LIKE family, we found that *PIP-LIKE6 (PIPL6*) expression is highly inducible to the exogenous application of chitin and infestation by the specialist aphid *Brevicoryne brassicae* (Vie et al., 2015). In the current study, we aimed at functionally characterizing the potential role of PIPL6 in the regulation of immunity in Arabidopsis. The kinetics of *PIPL6* expression in response to different classes of PAMPs and DAMP showed very rapid and transient induction (Fig. 1). A similar expression pattern in response to flg22 treatment was reported for *PIP1-3* peptides (Najafi et al., 2020) but the response of *PIPL6* to flg22 was much stronger. In addition, exogenous application of synthetic PIPL6 peptide elicited expression of immune marker genes in an RLK7-dependent manner, indicating that that RLK7 plays an important role as a receptor for PIPL6 initiated signalling in the regulation of immunity (Fig. 2).

Activation of the plant immune response is highly energy-demanding and has a negative impact on plant growth. For example, ectopic application or overexpression of PIP1 causes a significant reduction in Arabidopsis root growth (Hou et al., 2014). Analysis of root growth dynamics under both long and short day conditions showed that overexpression of PIPL6 had no visual effect on primary root growth and development (Supplementary Fig. 1). Profiling of the total transcriptome landscape three hours after exogenous application of the synthetic PIPL6 peptide revealed a massive reprogramming where the upregulated part is significantly enriched in detoxification- and immunity-related pathways including genes involved in JA, SA, ethylene, glucosinolate and camalexin biosynthesis and signalling (Table 1). Our data point towards a DAMP function for PIPL6. Similar to PAMPs, perception of DAMPs triggers several early signaling events, including the influx of calcium ions (Ca^2+^), activation of MAPK cascades, production of reactive oxygen species (ROS), and ethylene production (Zipfel, 2009). The early responses are followed by later events such as changes in gene expression patterns, and production of phytohormones and secondary metabolites. Furthermore, it has been shown that PEP1-initiated signalling in response to bacterial pathogens or flg22 is required to tailor the local and systemic responses. Localized application of PEP1 peptide co-activates the JA and SA branches in systemic leaves of Arabidopsis (Ross et al., 2014). Among the early events in PAMP and DAMP triggered signal transduction, phosphorylation of MAP-kinases play a critical role to convey the signal from the cell surface and through downstream modulators to the nucleus (Zhang and Klessig, 2001). Of the 20 MAPKs encoded by the Arabidopsis genome MAPK3, MAPK4, and MAPK6 are well studied for their critical function in signal transduction upon pathogen and elicitor perceptions ^43^. Recognition of phytopathogens and plant immune elicitors leads to rapid activation of the three Arabidopsis mitogen-activated protein kinases (MAPKs) MAPK3, MAPK4, and MAPK6. The importance of MAPK signaling cascades in plant–pathogen interactions is supported by the fact that many effectors that are secreted by pathogens target plant MAPK cascades to perturb PTI (Cui et al., 2010; Yu et al., 2020). Perception of flg22 activates two MAPK signalling pathways in parallel. The first module includes MKK4 and MKK5, which act redundantly to activate MAPK3 and MAPK6. The second cascade is composed of MEKK1, which activates MKK1 and MKK2 that redundantly phosphorylate MAPK4 (Asai et al., 2002). While it has been shown that PEP1 activates both modules (Gao et al., 2008), exogenous application of synthetic PIPL6 peptide activates only the MAPK3 and MPK6 module in an RLK7 dependent manner (Fig. 3a.). Activated MAPKs phosphorylate many substrates including different classes of TFs (Mao et al., 2011). Previous studies showed that upon Botrytis infection, WRKY33 serves as a direct substrate of MAPK3 and MAPK6 and that ectopic application of flg22 results in a fast induction of WRKY33 translation 1 h after treatment (Birkenbihl et al., 2017). Recently, it has been shown that elicitation of ROOT MERISTEM GROWTH FACTOR 7 (RGF7) peptide and its perception by RGF1-INSENSITIVE 4 and 5 (RGI4/5) receptors activates the MAPK6/3 and WRKY33 module to enhance Arabidopsis resistance against the bacterial pathogen *Pseudomonas syringae* (Wang et al., 2021). Similarly, PIPL6 is able to induce the translation of WRKY33 (Fig. 3b.). In addition, *PIPL6* expression is highly dependent on WRKY33 (Fig. 4b.). WRKY33 functions as a master regulator of plant immune response against necrotrophic fungal pathogens (Zheng et al., 2006; Mao et al., 2011; Najafi et al., 2020). WRKY33 exerts its role through the regulation of the hormonal balance between SA and JA/ET pathways, repression of ABA biosynthesis, and camalexin biosynthesis during infection. Arabidopsis resistance against necrotrophic pathogens is associated with a simultaneous activation of JA/ET and repression of SA and ABA signalling pathways. It has been shown that, 14 h after *Botrytis cinerea* infection, WRKY33 directly binds to the promoter regions of *JAZ1* and *JAZ5*, two repressors of MYC2, and represses their expression. Both JA and SA biosynthesis and signalling pathways were activated in *wrky33* mutant plants challenged by Botrytis in our study. Coactivation of JA and SA biosynthesis results in suppression of JA signalling by the SA pathway and results in triggering cell death, which acts in favor of necrotrophs (Spoel and Dong, 2008). ABA production during botrytis infection has a negative impact on plant immune response. WRKY33 directly controls ABA production by repression of *NCED3* and *NCED5* expression to restrict pathogen invasion (Liu et al., 2015). Furthermore, WRKY33 targets *CYP71A13* and *PAD3* which encode two major enzymes in camalexin biosynthesis in Arabidopsis and thereby positively regulates camalexin production. Measurements of SA, JA, ABA, and camalexin production in *PIPL6* transgenic lines 48 h after Botrytis infection revealed that the observed susceptibility in *pipl6* mutant lines is probably related to their compromised productions of JA and camalexin. The lower levels of glucosinolates in the *pipl6* mutants could also contribute to their increased susceptibility to Botrytis (Fig. 9b.). Indeed, secondary metabolites such as glucosinolates and their hydrolysis products play important roles in resistance of *Brassicaceae* to pathogens (Clay et al., 2009; Bednarek, 2012). Botrytis infection of Arabidopsis affects the expression of glucosinolate-related genes, such as *MYB28* and *MYB51*, causes changes in glucosinolate levels that differ among Botrytis isolates and plant genotypes, and differences in susceptibility to necrotrophic fungi of Arabidopsis mutants that are deficient in camalexin and/or glucosinolates has been shown (Kliebenstein et al., 2005; Clay et al., 2009; Xu et al., 2016; Castillo et al., 2019). It is worth noting that the applied isolate of Botrytis in this study is not able to discriminate the levels of resistance in *PIPL6:OX* background since the Wt plants exhibit resistant phenotype. This may explain the similar levels of quantified camalexin in Wt and *PIPL6:OX* lines (Fig. 8). Treatment of plants using spores from *Alternaria brassicae* was successfully discriminating a more resistant phenotype for *PIPL6:OX* lines than for Wt. This observation confirmed our hypothesis that PIPL6 functions as an amplifier of plant immune response against necrotrophic pathogens (Fig. 6 and Supplementary Fig. 2).

## Conclusion

Our findings revealed the presence of a loop where the perception of PIPL6 by RLK7 activates MPK6 and MPK3 upstream of WRKY33. In turn, activated WRKY33 regulates the expression of *PIPL6* itself and creates a positive feed-forward loop to amplify plant immune response through the regulation of plant phytohormones and secondary metabolites such as camalexin and glucosinolates (Fig. 10).

**Figure 10.**
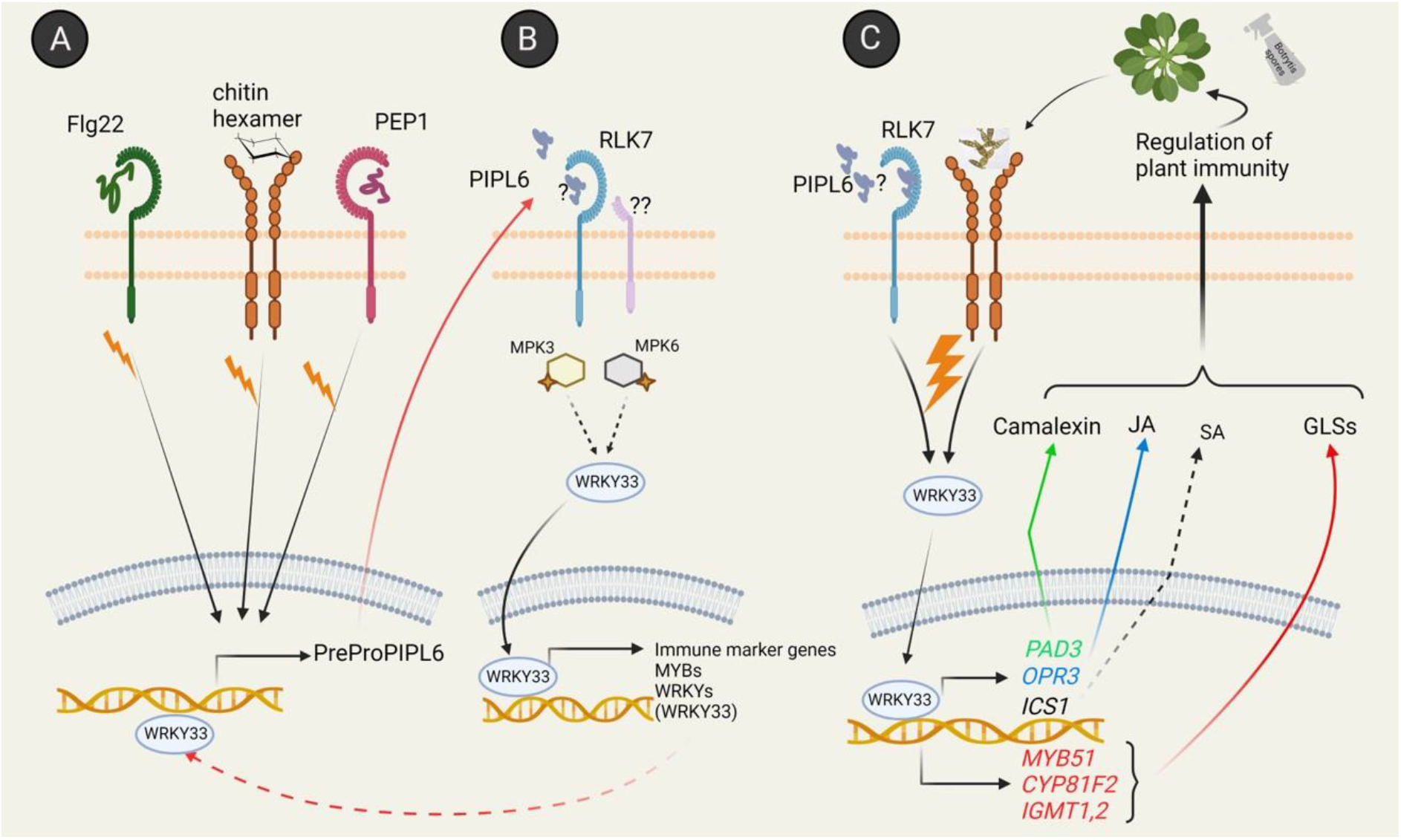
Proposed model for PIPL6 initiated signalling in the regulation of plant immunity. (A) *PIPL6* transcript level is induced by perception of PAMPs and DAMPs through their corresponding PRRs. B) Mature PIPL6 is most likely is perceived by RLK7 at plasma membrane and activates MPK6/3 and WRKY33 module and regulates expression of many genes involved in immunity. C) Upon infection by *B. cinerea*, activation of PIPL6 initiated signallin pathway, among the other known pathways, amplifies plant immune response through the regulation of JA, SA, camalexin, and glucosinolates biosynthesis to drive the plants ultimate response.

## Notes

### Competing Interest Statement

The authors have declared no competing interest.

## References

Ahuja I, Kissen R, Bones AM (2012) Phytoalexins in defense against pathogens. Trends in plant science 17: 73–90

Asai T, Tena G, Plotnikova J, Willmann MR, Chiu W-L, Gomez-Gomez L, Boller T, Ausubel FM, Sheen J (2002) MAP kinase signalling cascade in Arabidopsis innate immunity. Nature 415: 977–983

Bednarek P (2012) Chemical warfare or modulators of defence responses–the function of secondary metabolites in plant immunity. Current opinion in plant biology 15: 407–414

Berens ML, Berry HM, Mine A, Argueso CT, Tsuda K (2017) Evolution of hormone signaling networks in plant defense. Annual review of phytopathology 55: 401–425

Bindea G, Mlecnik B, Hackl H, Charoentong P, Tosolini M, Kirilovsky A, Fridman W-H, Pagès F, Trajanoski Z, Galon J (2009) ClueGO: a Cytoscape plug-in to decipher functionally grouped gene ontology and pathway annotation networks. Bioinformatics 25: 1091–1093

Birkenbihl RP, Diezel C, Somssich IE (2012) Arabidopsis WRKY33 is a key transcriptional regulator of hormonal and metabolic responses toward Botrytis cinerea infection. Plant physiology 159: 266–285

Birkenbihl RP, Kracher B, Roccaro M, Somssich IE (2017) Induced genome-wide binding of three Arabidopsis WRKY transcription factors during early MAMP-triggered immunity. The Plant Cell 29: 20–38

Bisceglia NG, Gravino M, Savatin DV (2015) Luminol-based assay for detection of immunity elicitor-induced hydrogen peroxide production in Arabidopsis thaliana leaves. Bio-protocol 5: e1685–e1685

Blažević I, Montaut S, Burčul F, Olsen CE, Burow M, Rollin P, Agerbirk N (2020) Glucosinolate structural diversity, identification, chemical synthesis and metabolism in plants. Phytochemistry 169: 112100

Boller T, Flury P (2012) Peptides as danger signals: MAMPs and DAMPs. In Plant signaling peptides. Springer, pp 163–181

Brown PD, Tokuhisa JG, Reichelt M, Gershenzon J (2003) Variation of glucosinolate accumulation among different organs and developmental stages of Arabidopsis thaliana. Phytochemistry 62: 471–481

Castillo N, Pastor V, Chávez Á, Arró M, Boronat A, Flors V, Ferrer A, Altabella T (2019) Inactivation of UDP-glucose sterol glucosyltransferases enhances Arabidopsis resistance to Botrytis cinerea. Frontiers in plant science 10: 1162

Clay NK, Adio AM, Denoux C, Jander G, Ausubel FM (2009) Glucosinolate metabolites required for an Arabidopsis innate immune response. Science 323: 95–101

Clough SJ, Bent AF (1998) Floral dip: a simplified method for Agrobacterium-mediated transformation of Arabidopsis thaliana. The plant journal 16: 735–743

Combest MM, Moroz N, Tanaka K, Rogan CJ, Anderson JC, Thura L, Rakotondrafara AM, Goyer A (2021) St PIP1, a PAMP-induced peptide in potato, elicits plant defenses and is associated with disease symptom severity in a compatible interaction with Potato virus Y. Journal of Experimental Botany 72: 4472–4488

Cui H, Wang Y, Xue L, Chu J, Yan C, Fu J, Chen M, Innes RW, Zhou J-M (2010) Pseudomonas syringae effector protein AvrB perturbs Arabidopsis hormone signaling by activating MAP kinase 4. Cell host & microbe 7: 164–175

De Mendiburu F (2014) Agricolae: statistical procedures for agricultural research. R package version 1: 1–4

Earley KW, Haag JR, Pontes O, Opper K, Juehne T, Song K, Pikaard CS (2006) Gateway-compatible vectors for plant functional genomics and proteomics. The Plant Journal 45: 616–629

Gao M, Liu J, Bi D, Zhang Z, Cheng F, Chen S, Zhang Y (2008) MEKK1, MKK1/MKK2 and MPK4 function together in a mitogen-activated protein kinase cascade to regulate innate immunity in plants. Cell research 18: 1190–1198

Gigolashvili T, Yatusevich R, Berger B, Müller C, Flügge UI (2007) The R2R3-MYB transcription factor HAG1/MYB28 is a regulator of methionine-derived glucosinolate biosynthesis in Arabidopsis thaliana. The plant journal 51: 247–261

Glazebrook J (2005) Contrasting mechanisms of defense against biotrophic and necrotrophic pathogens. Annual review of phytopathology 43: 205

Gómez-Gómez L, Boller T (2000) FLS2: an LRR receptor–like kinase involved in the perception of the bacterial elicitor flagellin in Arabidopsis. Molecular cell 5: 1003–1011

Graser G, Schneider B, Oldham NJ, Gershenzon J (2000) The methionine chain elongation pathway in the biosynthesis of glucosinolates in Eruca sativa (Brassicaceae). Archives of Biochemistry and Biophysics 378: 411–419

Hou S, Wang X, Chen D, Yang X, Wang M, Turrà D, Di Pietro A, Zhang W (2014) The secreted peptide PIP1 amplifies immunity through receptor-like kinase 7. PLoS pathogens 10: e1004331

Hu XY, Neill SJ, Cai WM, Tang ZC (2004) Induction of defence gene expression by oligogalacturonic acid requires increases in both cytosolic calcium and hydrogen peroxide in Arabidopsis thaliana. Cell research 14: 234–240

Hu Z, Zhang H, Shi K (2018) Plant peptides in plant defense responses. Plant signaling & behavior 13: e1475175

Huffaker A, Pearce G, Ryan CA (2006) An endogenous peptide signal in Arabidopsis activates components of the innate immune response. Proceedings of the National Academy of Sciences 103: 10098–10103

Jones JD, Dangl JL (2006) The plant immune system. nature 444: 323–329

Kissen R, Eberl F, Winge P, Uleberg E, Martinussen I, Bones AM (2016) Effect of growth temperature on glucosinolate profiles in Arabidopsis thaliana accessions. Phytochemistry 130: 106–118

Kliebenstein DJ, Rowe HC, Denby KJ (2005) Secondary metabolites influence Arabidopsis/Botrytis interactions: variation in host production and pathogen sensitivity. The Plant Journal 44: 25–36

Krol E, Mentzel T, Chinchilla D, Boller T, Felix G, Kemmerling B, Postel S, Arents M, Jeworutzki E, Al-Rasheid KA (2010) Perception of the Arabidopsis danger signal peptide 1 involves the pattern recognition receptor AtPEPR1 and its close homologue AtPEPR2. Journal of Biological Chemistry 285: 13471–13479

Langmead B, Salzberg SL (2012) Fast gapped-read alignment with Bowtie 2. Nature methods 9: 357–359

Ligterink W, Kroj T, Nieden Uz, Hirt H, Scheel D (1997) Receptor-mediated activation of a MAP kinase in pathogen defense of plants. Science 276: 2054–2057

Liu S, Kracher B, Ziegler J, Birkenbihl RP, Somssich IE (2015) Negative regulation of ABA signaling by WRKY33 is critical for Arabidopsis immunity towards Botrytis cinerea 2100. Elife 4: e07295

Liu Z, Wu Y, Yang F, Zhang Y, Chen S, Xie Q, Tian X, Zhou J-M (2013) BIK1 interacts with PEPRs to mediate ethylene-induced immunity. Proceedings of the National Academy of Sciences 110: 6205–6210

Luna E, Pastor V, Robert J, Flors V, Mauch-Mani B, Ton J (2011) Callose deposition: a multifaceted plant defense response. Molecular Plant-Microbe Interactions 24: 183–193

Maleck K, Levine A, Eulgem T, Morgan A, Schmid J, Lawton KA, Dangl JL, Dietrich RA (2000) The transcriptome of Arabidopsis thaliana during systemic acquired resistance. Nature genetics 26: 403–410

Mao G, Meng X, Liu Y, Zheng Z, Chen Z, Zhang S (2011) Phosphorylation of a WRKY transcription factor by two pathogen-responsive MAPKs drives phytoalexin biosynthesis in Arabidopsis. The Plant Cell 23: 1639–1653

Miller G, Schlauch K, Tam R, Cortes D, Torres MA, Shulaev V, Dangl JL, Mittler R (2009) The plant NADPH oxidase RBOHD mediates rapid systemic signaling in response to diverse stimuli. Science signaling 2: ra45–ra45

Miya A, Albert P, Shinya T, Desaki Y, Ichimura K, Shirasu K, Narusaka Y, Kawakami N, Kaku H, Shibuya N (2007) CERK1, a LysM receptor kinase, is essential for chitin elicitor signaling in Arabidopsis. Proceedings of the National Academy of Sciences 104: 19613–19618

Najafi J, Brembu T, Vie AK, Viste R, Winge P, Somssich IE, Bones AM (2020) PAMP-induced secreted peptide 3 modulates immunity in Arabidopsis. Journal of Experimental Botany 71: 850–864

Robinson MD, McCarthy DJ, Smyth GK (2010) edgeR: a Bioconductor package for differential expression analysis of digital gene expression data. bioinformatics 26: 139–140

Ross A, Yamada K, Hiruma K, Yamashita-Yamada M, Lu X, Takano Y, Tsuda K, Saijo Y (2014) The Arabidopsis PEPR pathway couples local and systemic plant immunity. The EMBO journal 33: 62–75

Salem MA, Yoshida T, Perez de Souza L, Alseekh S, Bajdzienko K, Fernie AR, Giavalisco P (2020) An improved extraction method enables the comprehensive analysis of lipids, proteins, metabolites and phytohormones from a single sample of leaf tissue under water-deficit stress. The Plant Journal 103: 1614–1632

Spoel SH, Dong X (2008) Making sense of hormone crosstalk during plant immune responses. Cell host & microbe 3: 348–351

Vie AK, Najafi J, Liu B, Winge P, Butenko MA, Hornslien KS, Kumpf R, Aalen RB, Bones AM, Brembu T (2015) The IDA/IDA-LIKE and PIP/PIP-LIKE gene families in Arabidopsis: phylogenetic relationship, expression patterns, and transcriptional effect of the PIPL3 peptide. Journal of Experimental Botany 66: 5351–5365

Wang X, Zhang N, Zhang L, He Y, Cai C, Zhou J, Li J, Meng X (2021) Perception of the pathogen-induced peptide RGF7 by the receptor-like kinases RGI4 and RGI5 triggers innate immunity in Arabidopsis thaliana. New Phytologist 230: 1110–1125

Xu J, Meng J, Meng X, Zhao Y, Liu J, Sun T, Liu Y, Wang Q, Zhang S (2016) Pathogen-responsive MPK3 and MPK6 reprogram the biosynthesis of indole glucosinolates and their derivatives in Arabidopsis immunity. The Plant Cell 28: 1144–1162

Yamaguchi Y, Huffaker A, Bryan AC, Tax FE, Ryan CA (2010) PEPR2 is a second receptor for the Pep1 and Pep2 peptides and contributes to defense responses in Arabidopsis. The Plant Cell 22: 508–522

Yu G, Xian L, Xue H, Yu W, Rufian JS, Sang Y, Morcillo RJ, Wang Y, Macho AP (2020) A bacterial effector protein prevents MAPK-mediated phosphorylation of SGT1 to suppress plant immunity. PLoS pathogens 16: e1008933

Zhang S, Klessig DF (2001) MAPK cascades in plant defense signaling. Trends in plant science 6: 520–527

Zheng Z, Qamar SA, Chen Z, Mengiste T (2006) Arabidopsis WRKY33 transcription factor is required for resistance to necrotrophic fungal pathogens. The Plant Journal 48: 592–605

Zipfel C (2009) Early molecular events in PAMP-triggered immunity. Current opinion in plant biology 12: 414–420

Zipfel C, Kunze G, Chinchilla D, Caniard A, Jones JD, Boller T, Felix G (2006) Perception of the bacterial PAMP EF-Tu by the receptor EFR restricts Agrobacterium-mediated transformation. Cell 125: 749–760

