## Supplementary figures for "PAMP-Induced secreted Peptide-Like 6 (PIPL6) functions as an amplifier of plant immune response through RLK7 and WRKY33 module"

Figure S1:

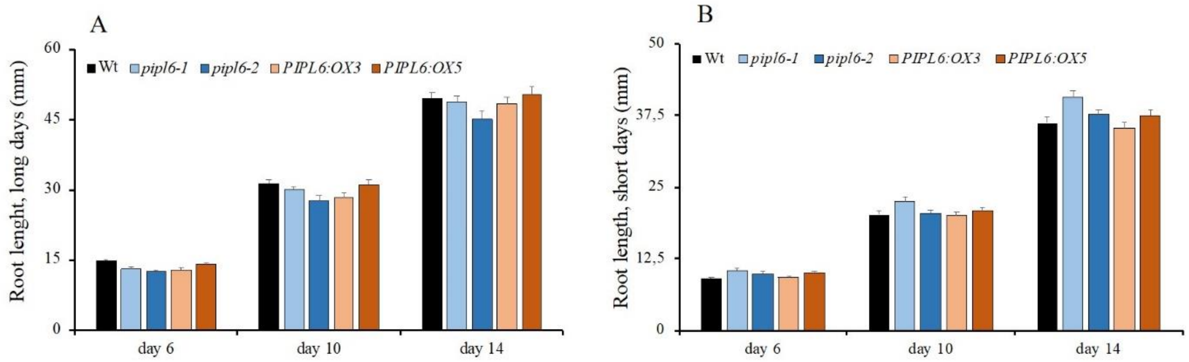


**Figure S1:** Overexpression or downregulation of PIPL6 has not a profound effect on root growth. Wt and *PIPL6* transgenic lines were grown in 1/2MS plates supplemented by 1% sucrose in a vertical position under both long days A) (16h light / 8h dark) and B) short days (10h light / 14h dark) regimes and the root growth dynamics were measured at mentioned time point. Bars and error bars are calculated from n>50 seedlings.

**Figure S2**

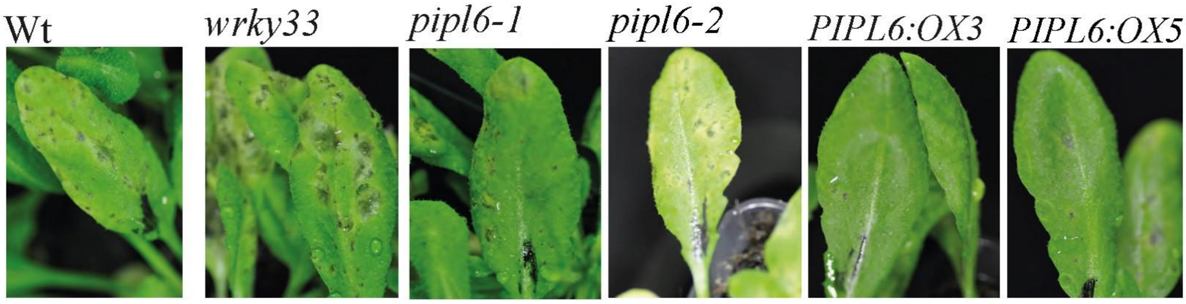


**Figure S2:** Overexpression of *PIPL6* enhances plant resistance against the *Alternaria brassicae* infection. Five-weeks-old Wt and transgenic lines were sprayed by *Alternaria brassicae* spores and plants were photographed five days after inoculation.

**Figure S3:**

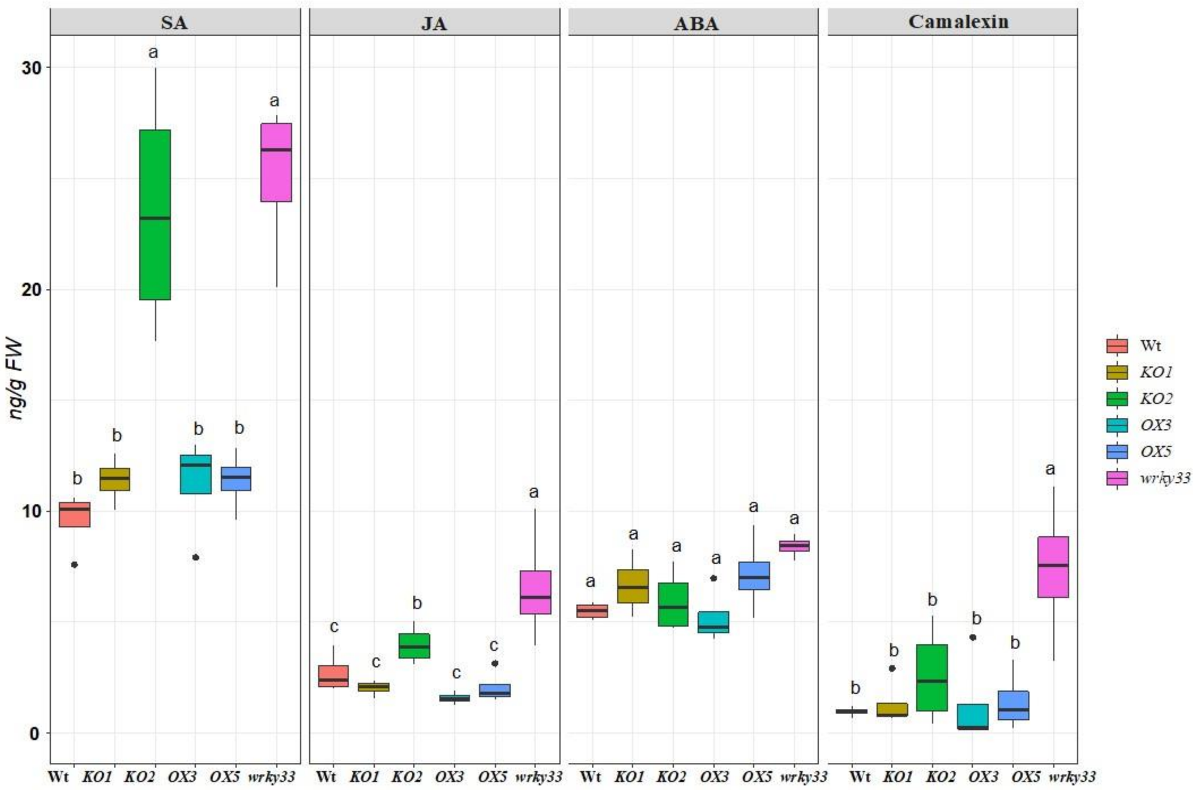


**Figure S3:** Analysis of hormonal content after mock treatment. Five-weeks-old plants were treated by Vogel buffer and rosette leaves were harvested 48hpi for hormonal extraction and quantification. Different letters indicate statistically significant differences calculated by ANOVA followed by Tukey multiple mean comparison method (n = 4, *p-value* $< 0.05$).

**Figure S4:**

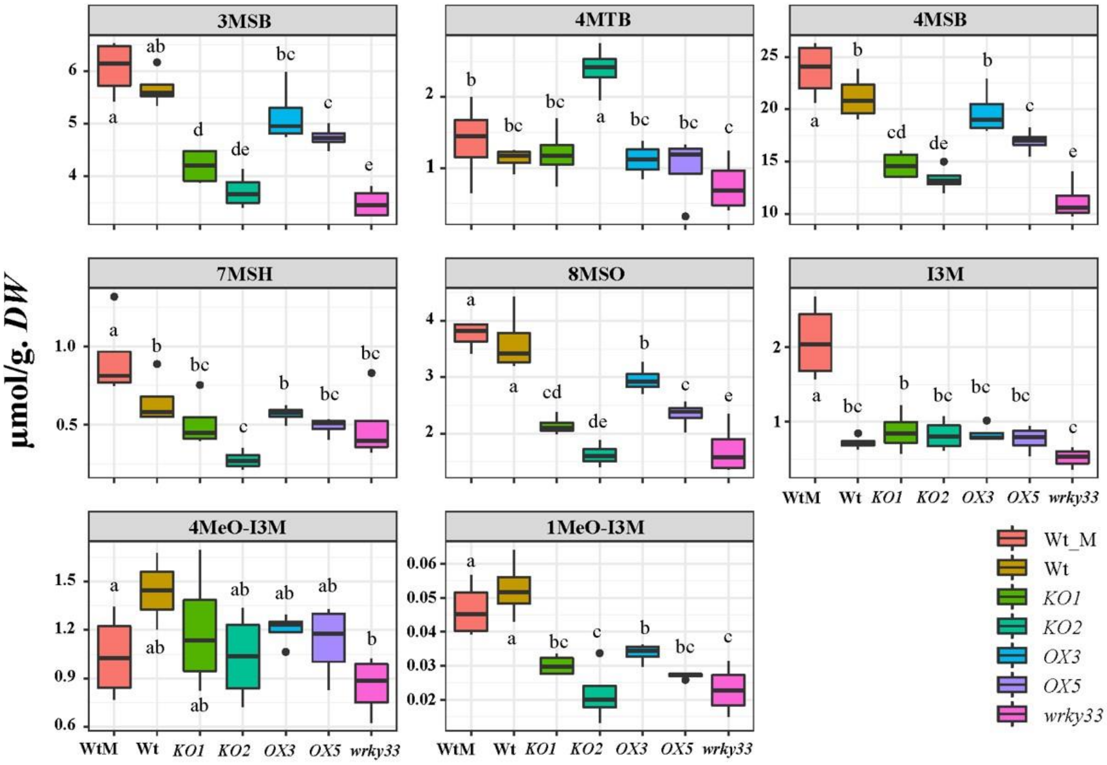


**Figure S4:** Detailed glucosinolates profile of studied genotypes challenged by *Botrytis cinerea* spores. Five-weeks-old plants were challenged by Botrytis spores and rosette leaves were harvested 48hpi for glucosinolates extraction, quantification, and detection of different class of glucosinolates. Different letters indicate statistically significant differences calculated by ANOVA followed by Tukey multiple mean comparison method (n = 4, *p-value* < 0.05).
